# The NeuroHab: A Low-Cost, Integrated System for Investigation of Neural Correlates of Behaviors

**DOI:** 10.64898/2026.08.09.743755

**Authors:** Samuel Crouse, Williams Johnston, Qian-Quan Sun

## Abstract

The development of a new integrated operant system was driven by two challenges in behavioral neuroscience: the high cost and technical complexity of commercial rigs, and their limited adaptability across experiments. We developed the NeuroHab, an integrated behavioral arena for high-fidelity operant conditioning and automated data collection in a single unified system. Food and water reward, conditioned-stimulus presentation, and event recording are tied together programmatically with easy-to-install open-source code to facilitate throughput and reproducibility. All behavioral events are processed by internal microcontrollers and logged with <1 ms latency (typical range 56–728 μs). This precise timing is critical for integrating the system with two-photon imaging and electrophysiology, enabling real-time alignment of behavior with brain activity. The NeuroHab uses solenoid-actuated, capacitive-sensing Lickports that let an untethered mouse drink from an automated port, and delivers food via the Kravitz Lab FED3. Conditioned stimuli are presented by dedicated buzzer/LED modules. A central controller (the Core) coordinates all modules and logs event timestamps using TTL-low signaling between two microcontrollers, at a maximum recording rate of 16.67 Hz for single-pulse events. We have deployed the NeuroHab in over 50 behavior trials and over 20 sessions alongside a Mini two-photon microscope. At approximately $1,400, easily modified, and compatible with existing analysis tools, the NeuroHab lowers barriers to multimodal behavioral neuroscience.

**Significance Statement:** The study of how neural activity gives rise to behavior depends on operant systems that are both temporally precise and affordable, yet commercial rigs are costly and difficult to adapt across experiments. We introduce the NeuroHab, an integrated, open-source operant platform that unifies reward delivery, conditioned-stimulus presentation, and event logging with sub-millisecond timing (typical latency 56–728 μs). Built for approximately $1,400, the system forwards all behavioral timestamps to external acquisition hardware, enabling millisecond-scale alignment of behavior with two-photon imaging and electrophysiology. By lowering the cost and technical barriers to synchronized behavioral and neural recording, the NeuroHab makes multimodal, reproducible operant neuroscience accessible to a broad range of laboratories and adaptable to diverse experimental paradigms.

## 1. Introduction

Studying behavior in mice is essential for determining the downstream effects of drug administration or neurological disorders (Crawley, 2012). When combined with imaging technologies, behavioral assays enable researchers to understand the functions of various brain regions. However, the experiments designed to determine such effects are often elaborate and expensive (Akam et al., 2022; Kapanaiah et al., 2021). It is therefore important that any rig used for conducting these experiments be accurate, modifiable, and facilitate repeatability while being suited to the specific experimental design.

Such rigs are not generally available commercially, forcing most researchers to build custom setups for each experiment. This approach consumes considerable time and resources. Moreover, complex or precision setups may be unreliable or require extensive testing periods before integration. This leads to redundant engineering effort and deadweight loss through duplication of common infrastructure including water/drug delivery, food delivery, conditioned stimuli, video recording, pipeline integration, and event logging. Additionally, the logging systems for bespoke solutions are often untested, imprecise, or difficult to integrate with existing workflows or data collection methods (Akam et al., 2022; Kapanaiah et al., 2021).

To address these challenges in our own research, we developed the NeuroHab as a modular, open-source platform with integrated operant design, precise temporal synchronization capabilities, and support for adaptable experimental paradigms. Our design philosophy centered on three principles: standardization of common behavioral components to maximize reproducibility, modular architecture analogous to object-oriented programming that breaks complex workflows into manageable parts, and unified control through the NeuroHab Core programming and logging system. This approach eliminates redundant design costs and enables flexible combinations of behavioral experiments.

The NeuroHab achieves <1 ms behavior event to logging latency (typical range: 56–728 μs) (Figures 2, 8) across all Core modules, enabling millisecond-scale temporal alignment with neural recording modalities including two-photon (Zong et al., 2017) calcium imaging, electrophysiology, and fiber-photometry (Figures 10a-11, 14). Unlike commercial alternatives that often require proprietary software and offer limited customization, the NeuroHab uses open-source code that researchers can freely modify to suit their experimental needs. The complete system can be assembled for approximately $1,400, approximately 7 times less expensive than comparable commercial operant conditioning systems (Med Associates Inc., 2026). The system has been validated across over 50 behavior trials and successfully integrated with Mini two-photon equipment (Figure 1).

**Fig. 1.**
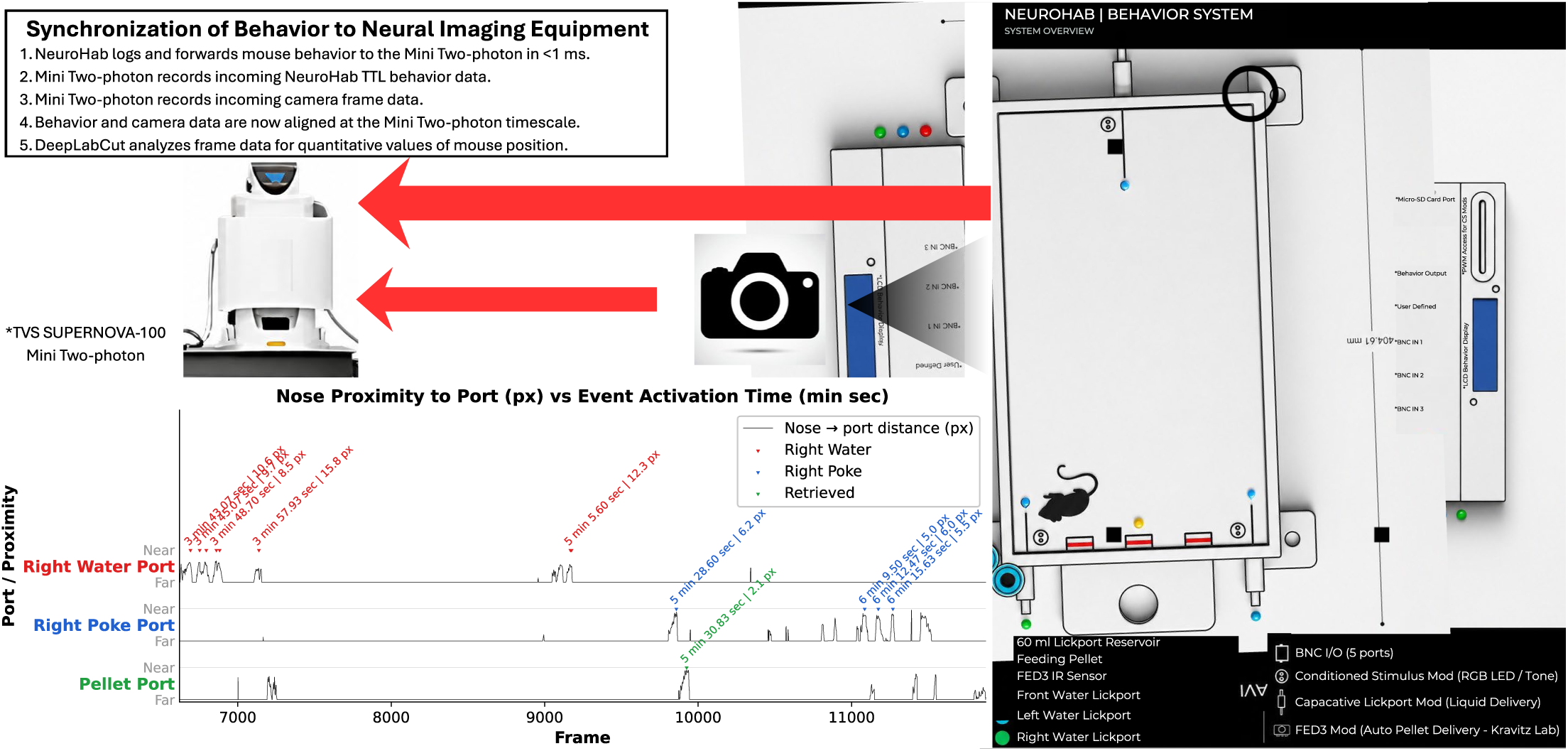
NeuroHab system illustration and synchronization with neural equipment experiment. NeuroHab directed and logged mouse behavior was recorded via Mini Two-photon (M2P) directed cameras. Mouse behavior events are forwarded in real time over the BNC connection from the NeuroHab to the M2P. A DeepLabCut (Mathis et al., 2018) model was trained on the captured frame data and a quantitative analysis of mouse position as distance from various ports was produced as seen in Figure 11. The behavior activation events for each port are overlaid on top of this data to illustrate that as the mouse approaches a port, an activation may have occurred. These data were synchronized in real time on the M2P system, <1 ms, and are compatible with future miniaturized two-photon microscopy imaging. Additionally, similar approaches with electrophysiology or fiber-photometry equipment are feasible (Figure 14). The NeuroHab further supports standalone experiments for behavior only paradigms with onboard logging and control structures as flagship features.

NeuroHab builds upon a growing ecosystem of open-source behavioral control systems. Python-based frameworks such as pyControl (Akam et al., 2022) and low-cost 5-choice operant box designs (Kapanaiah et al., 2021) have substantially lowered the cost and increased the transparency of rodent behavioral hardware. More recently, FreiBox (De La Crompe et al., 2023) has demonstrated an open-source Arduino Mega 2560-based behavioral setup for freely moving mice with submillisecond lick tracking and integration with 1-photon calcium imaging via the Miniscope V4, further advancing the accessibility of combined behavioral-neural experiments. NeuroHab is distinguished from these systems along several principal axes. First, in timing precision: NeuroHab provides a demonstrated sub-millisecond (<1 ms) behavior-to-log latency (typical range 56–728 μs), quantified through direct timing measurements (Figures 2, 8, 12, 13). Notably, this latency is achieved on the Arduino Mega 2560’s 16 MHz ATmega2560 processor, a slower and more widely accessible platform than the 168 MHz ARM Cortex-M4 pyboard used by pyControl (Akam et al., 2022). Second, in experimental context: NeuroHab is designed for home-cage and free-moving rodent paradigms, whereas many existing systems are optimized primarily for traditional operant chambers or head-fixed preparations. While FreiBox (De La Crompe et al., 2023) also supports freely moving paradigms, NeuroHab’s distinguishing architectural feature is its dual-microcontroller design, which separates behavioral control (Arduino) from event logging (ESP32) at the hardware level, whereas FreiBox employs a single Arduino for both functions and requires an external logger for <1 ms logging. Third, in neural integration: NeuroHab provides native BNC-based interfaces for direct synchronization with two-photon imaging, electrophysiology, and fiber-photometry (Figure 14) hardware, reducing reliance on custom synchronization hardware; FreiBox (De La Crompe et al., 2023) has demonstrated 1-photon imaging integration, and NeuroHab extends this to two-photon microscopy with a dedicated TTL pulse-count encoding scheme for event synchronization. In this way, NeuroHab advances beyond prior open-source tools not only in cost but also in timing fidelity, modularity, and integration with neural recording equipment.

**Fig. 2.**
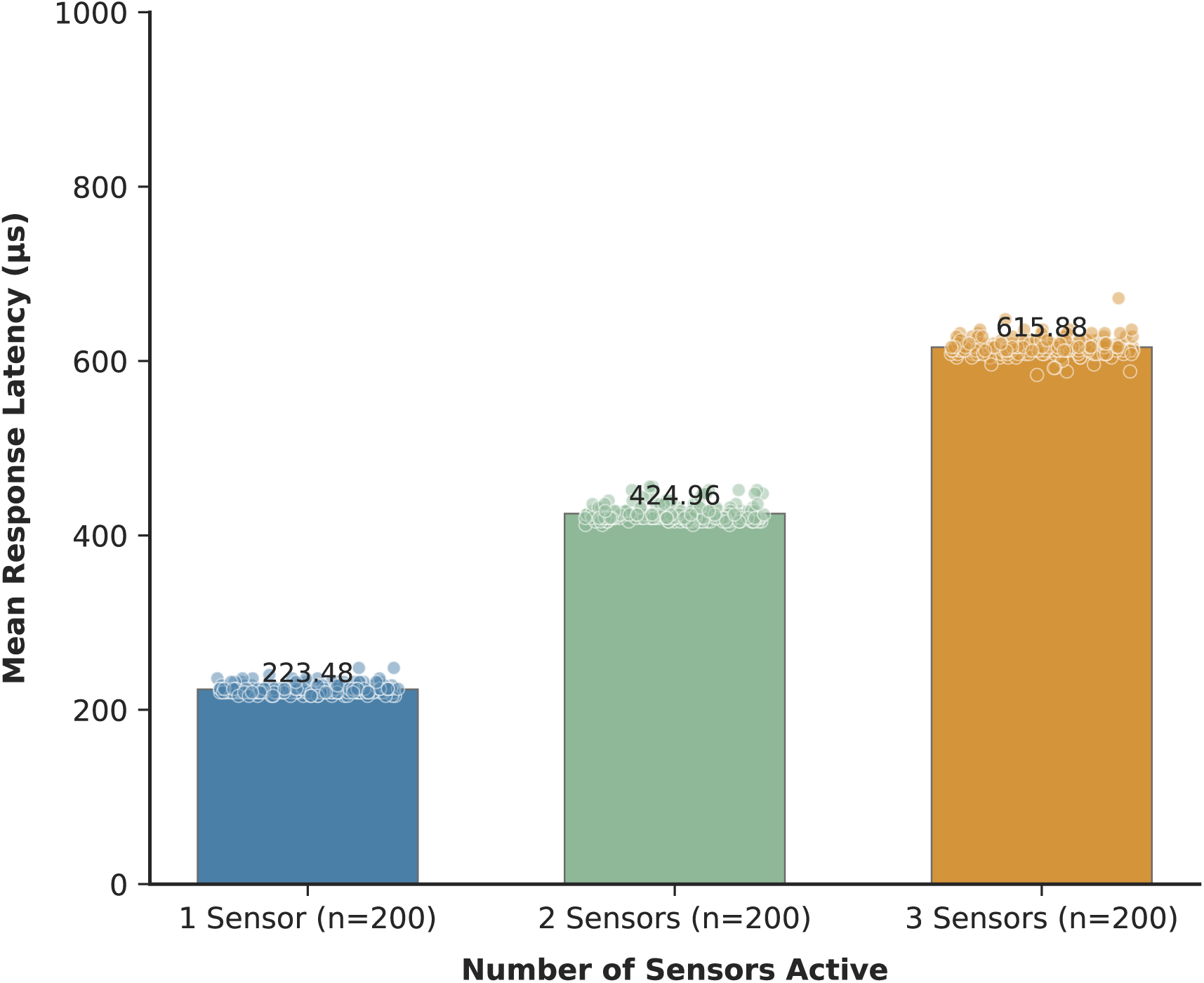
Lickport response latency across single, dual, and triple channel configurations. Bars represent mean latency (μs); Lickport capacitive sensors exhibit mean response latencies of 223.48 μs (SD = 5.49 μs, n = 200), 424.96 μs (SD = 8.84 μs, n = 200), and 615.88 μs (SD = 9.90 μs, n = 200) with 1, 2, and 3 Lickport sensors active.

**Fig. 3.**
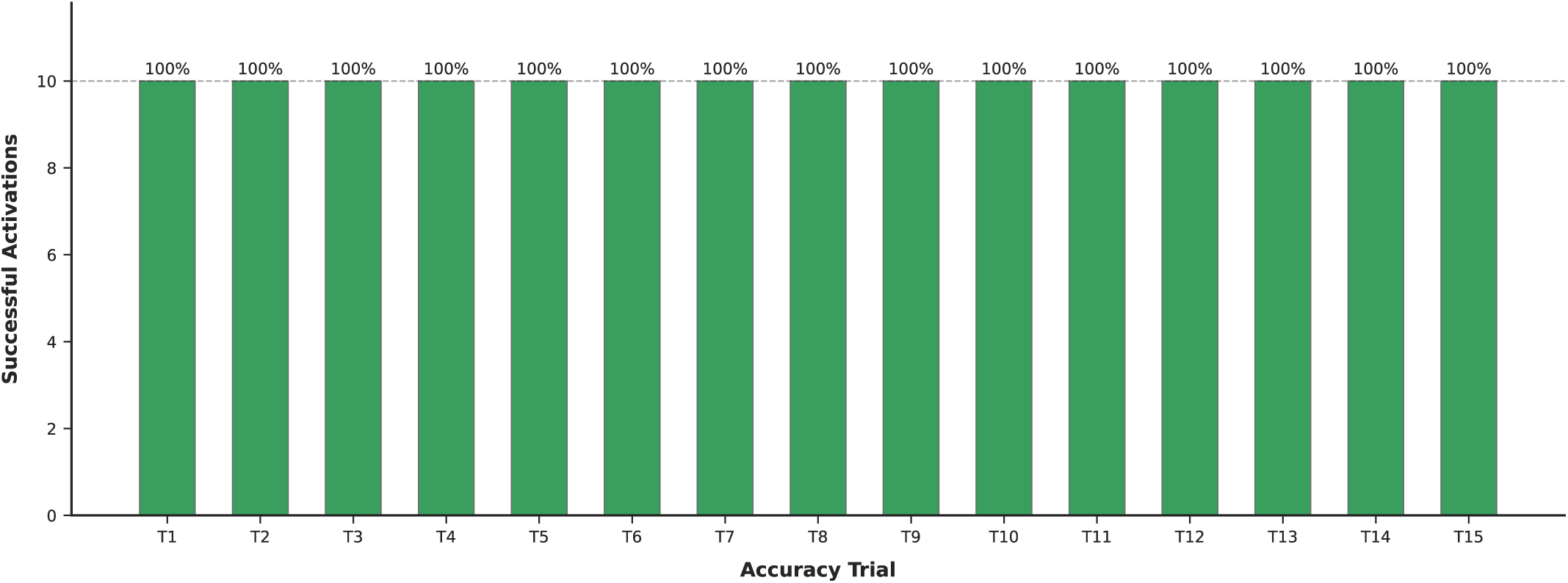
Lickport accuracy validation over 15 trials x 10 samples (n=150), 150/150 activations (100% overall). Tested over activation conditions (touch-and-release, touch-and-hold, rapid touch-and-release) on dry and wet surfaces.

**Fig. 4.**
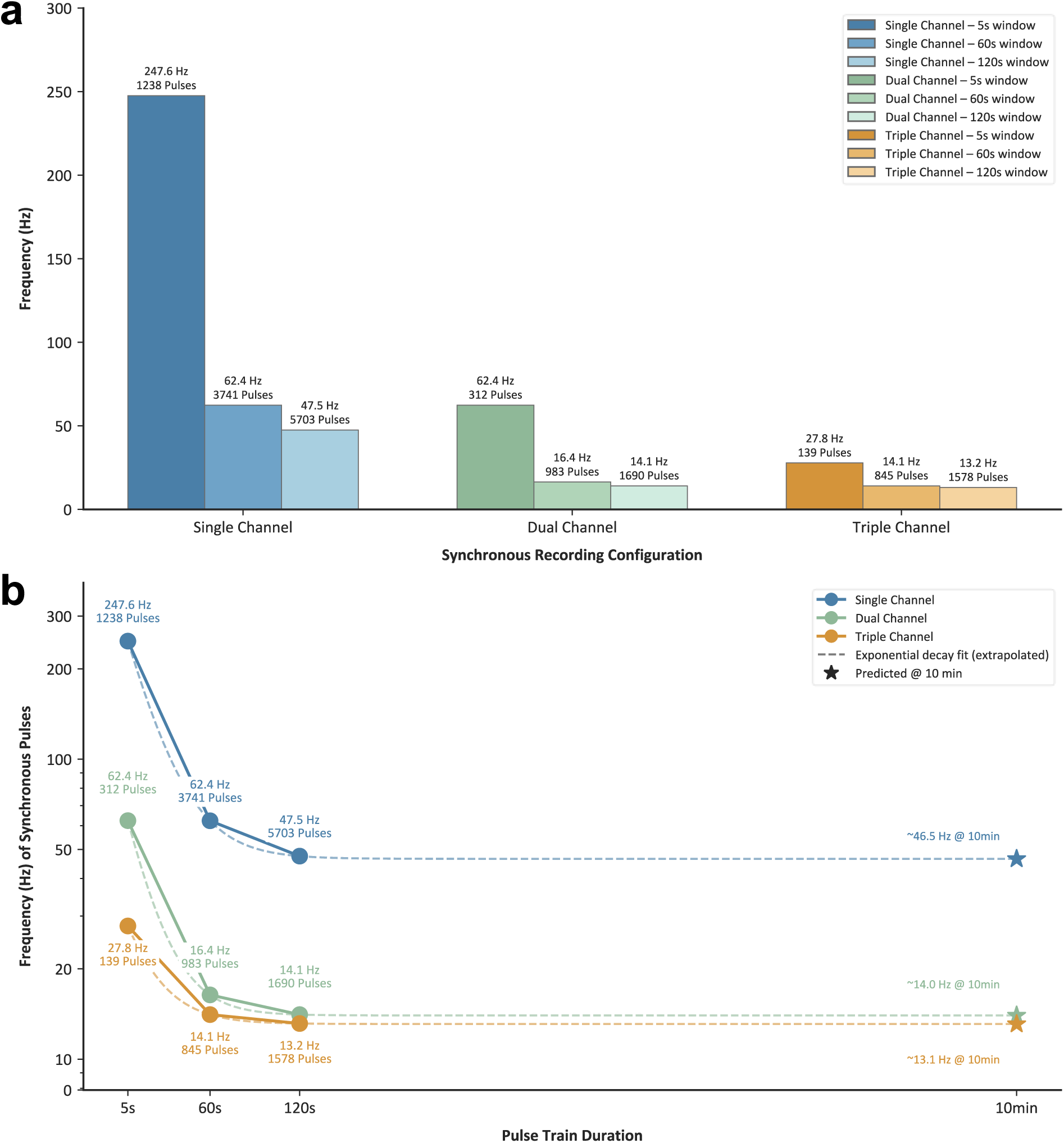
BNC input frequency validation of sustained synchronous pulses across channel configurations. **a.** BNC frequency maximums for single, dual, and triple channel recording with synchronous pulse trains. Values confirmed over three trials each at 100% detection rate. **b.** Maximum synchronous BNC recording frequencies by channel with predicted maximum frequencies at 10 minutes. Figure 4a single, dual, and triple channel values extrapolated out to predicted values at 10 minutes.

**Fig. 5.**
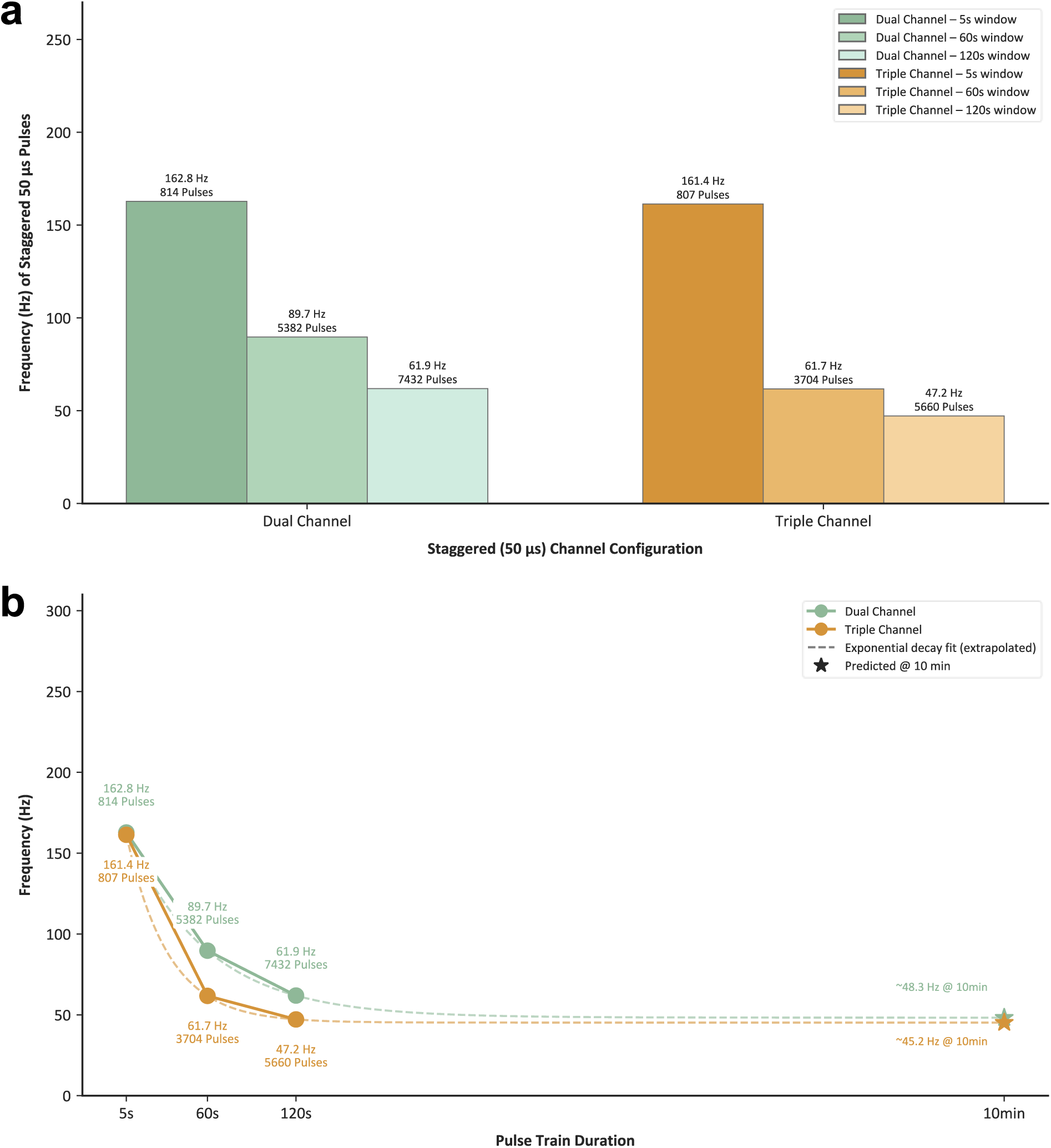
BNC input frequency validation of sustained staggered (50µs) pulses across dual and triple channel configurations. **a.** BNC frequency maximums for dual channel and triple channel recording with staggered pulse trains. Values confirmed over three trials each at 100% detection rate. **b.** Figure 5a dual, and triple channel values extrapolated out to predicted values at 10 minutes.

**Fig. 6.**
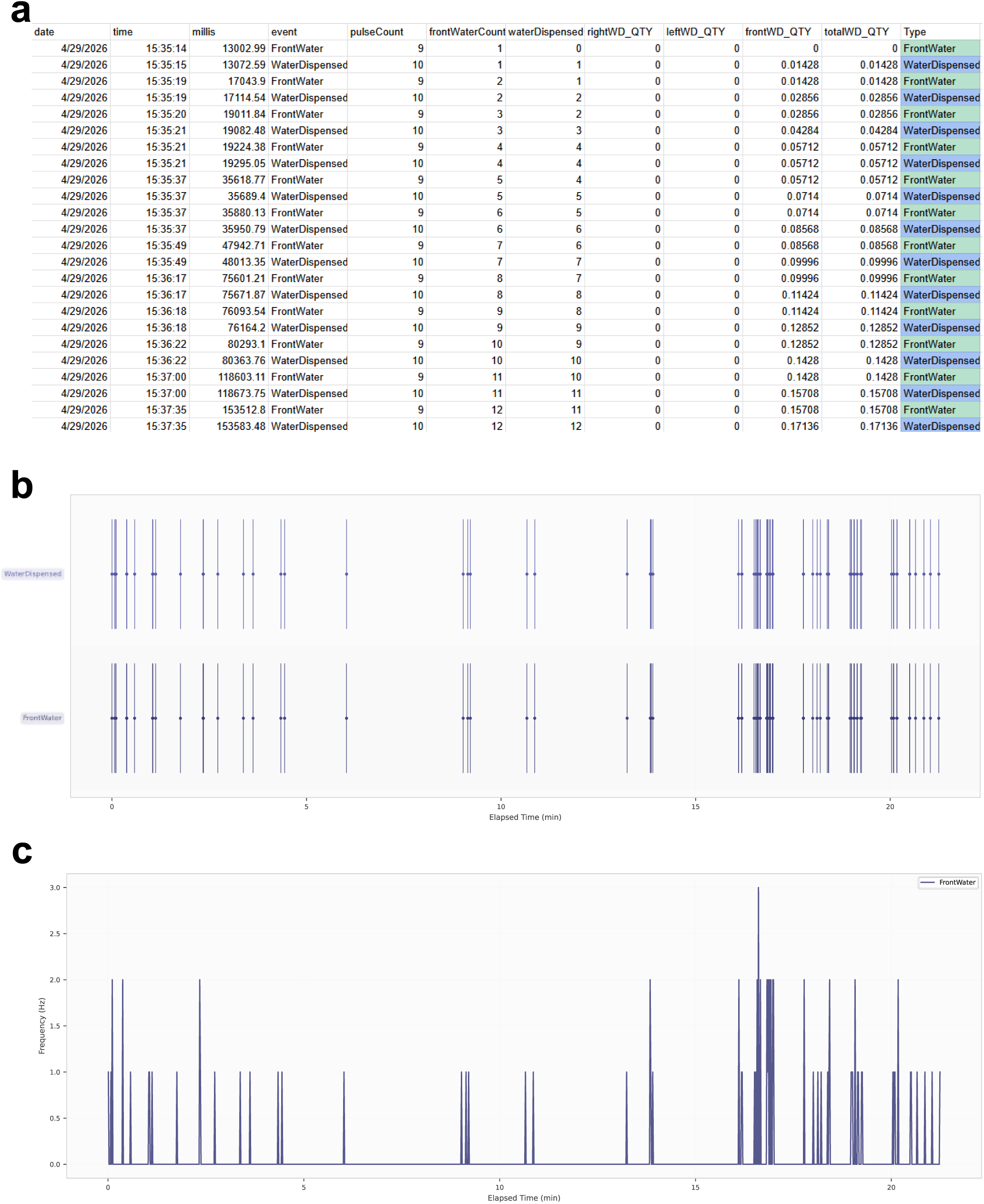
Head-fixed licking data analysis subset. **a.** Representative NeuroHab.csv output. **b.** Event raster for “FrontWater” and “WaterDispensed” events. **c.** “FrontWater” event frequencies over the trial smoothed at 1 second. Maximum instantaneous frequency recorded at 7.06 Hz with the average for bouts under 1 second at 2.96 Hz.

**Fig. 7.**
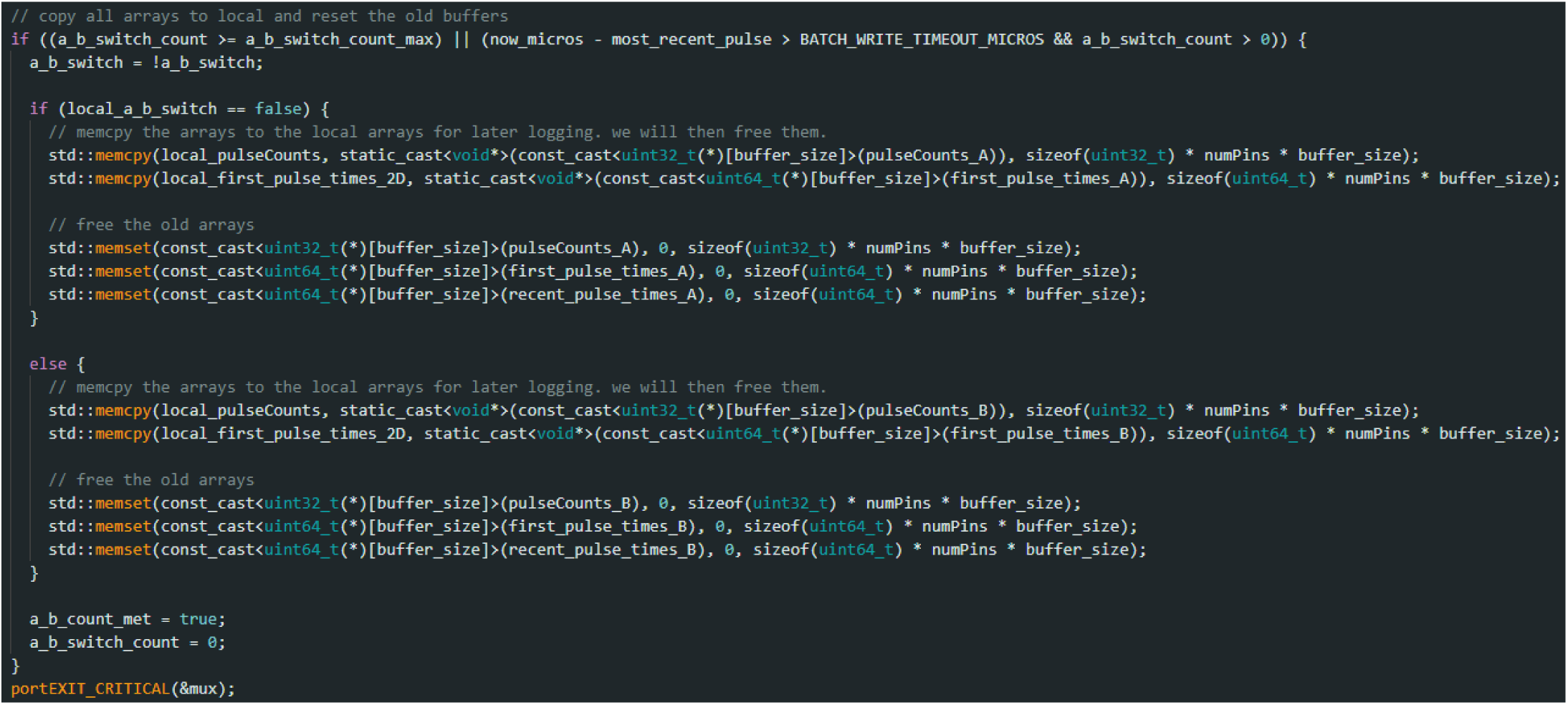
ESP32 buffer switching architecture within critical section. memcpy and memset for buffer switching and reallocation. Representative of architecture decisions for logging structure.

**Fig. 8.**
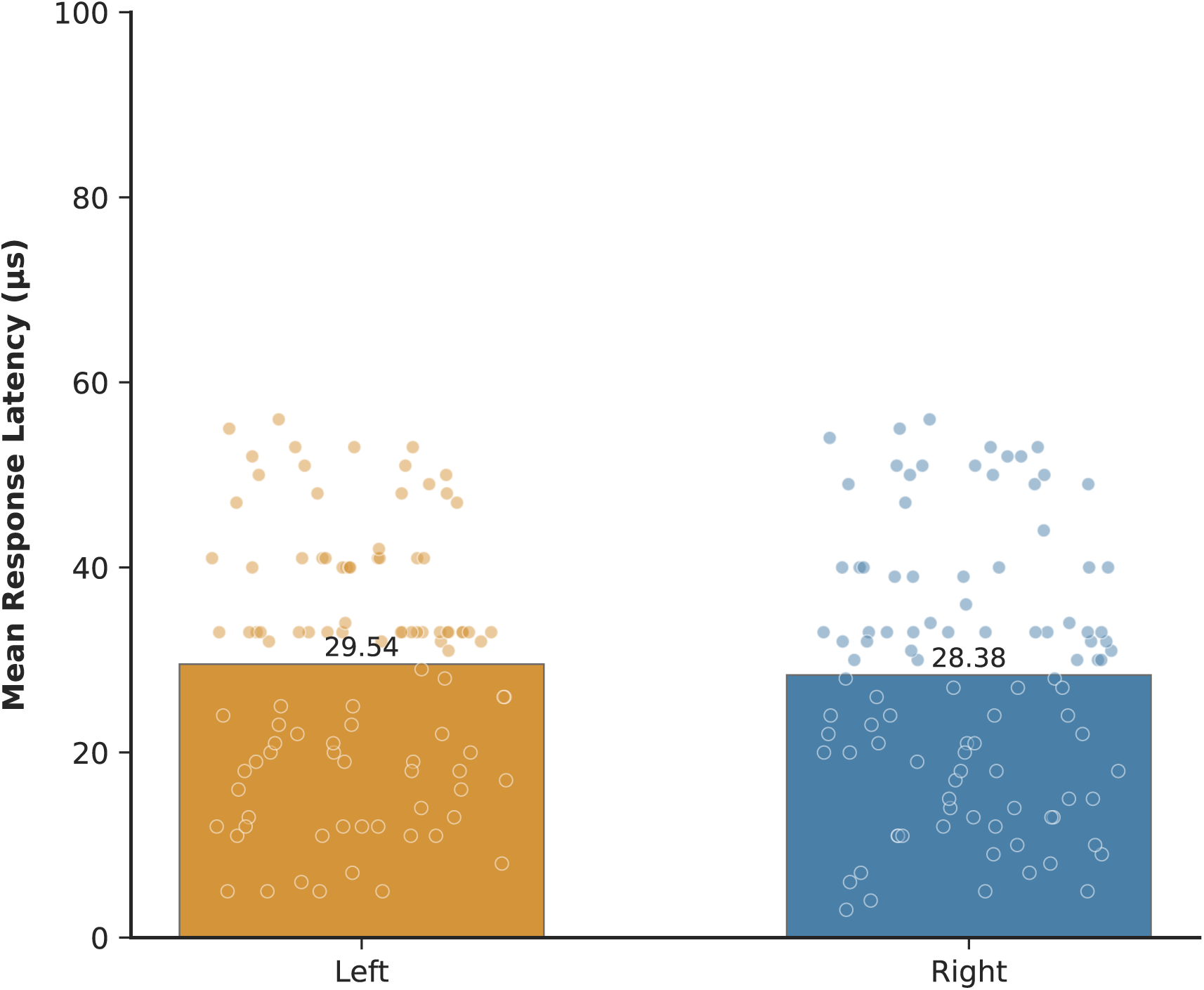
FED3 IR sensor response latency following NeuroHab integration. Mean latency = 28.96 μs ± 14.13 μs, n = 100 samples per port. Note: latency measured post-source code modification; stock FED3 latency is >1ms.

**Fig. 9.**
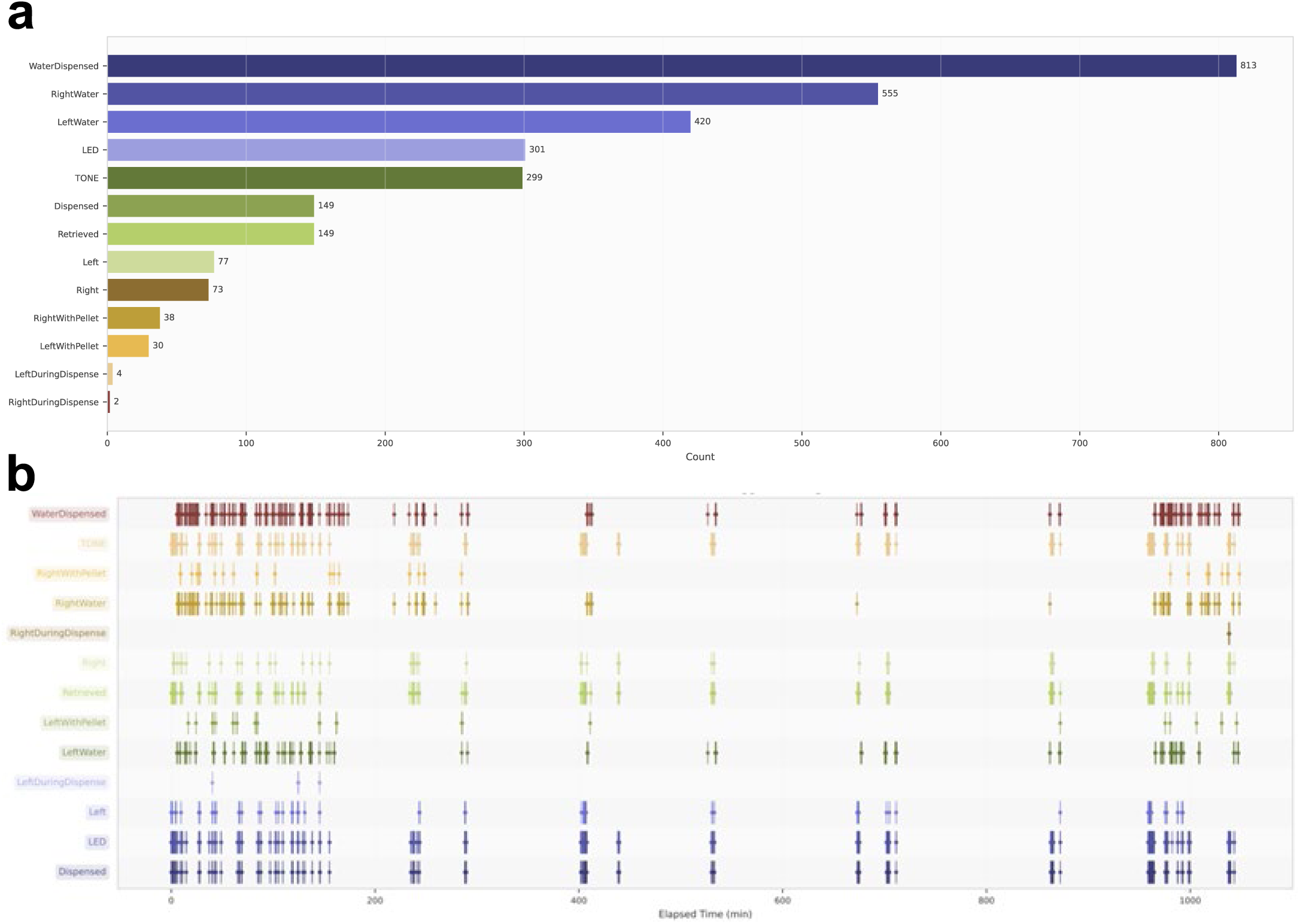
16-hour NeuroHab recording representative event plots. **a.** Event counts across a continuous 16-hour session showing water delivery, food retrieval, and nose poke events. **b.** Event raster for timing and alignment of events.

**Fig. 10.**
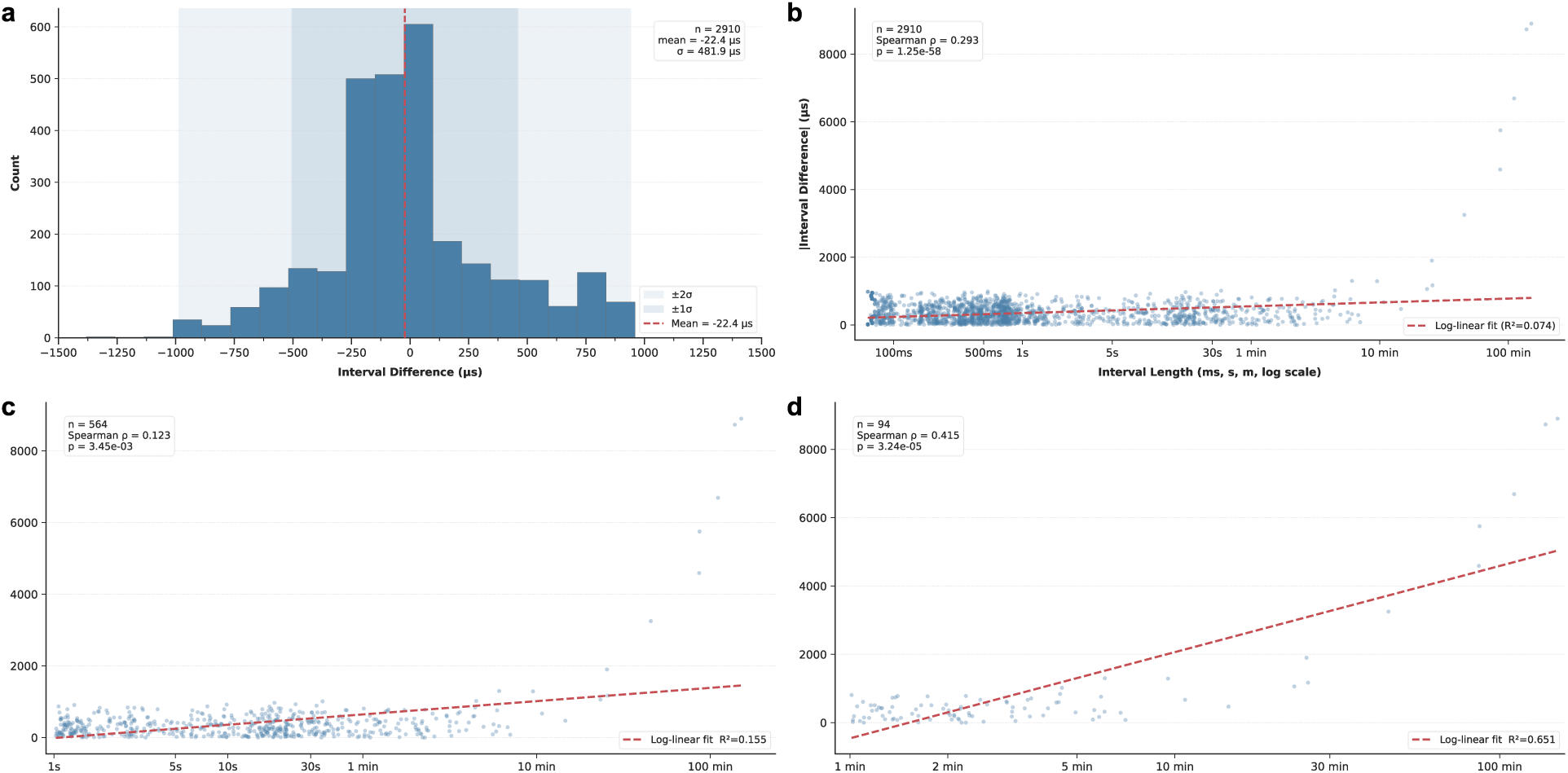
Microsecond interval distance distribution characterizes the temporal synchronization between the NeuroHab BNC and M2P recording systems. As time between events increases, clock drift accounts for more syncing discrepancy variance. **a.** The distribution of interval differences across all recorded pulse pairs, centered near zero (mean=−22.4 µs, n=2910, SD=481.9 µs) with spread bounded by ±1000 µs, a pattern consistent with M2P’s 1 ms timestamp resolution acting as a quantization noise floor rather than true timing error. **b.** Total event plot of absolute interval difference against interval length. **b – d** plot absolute interval difference against interval length across progressively stricter cutoffs (all data, ≥1 s, ≥1 min), illustrating how explained variance increases from **b:** R²=0.074, **c:** R²=0.155, to **d:** R²=0.651 as quantization noise is removed from shorter intervals, revealing the underlying clock drift signal that dominates at longer inter-event durations.

**Fig. 11.**
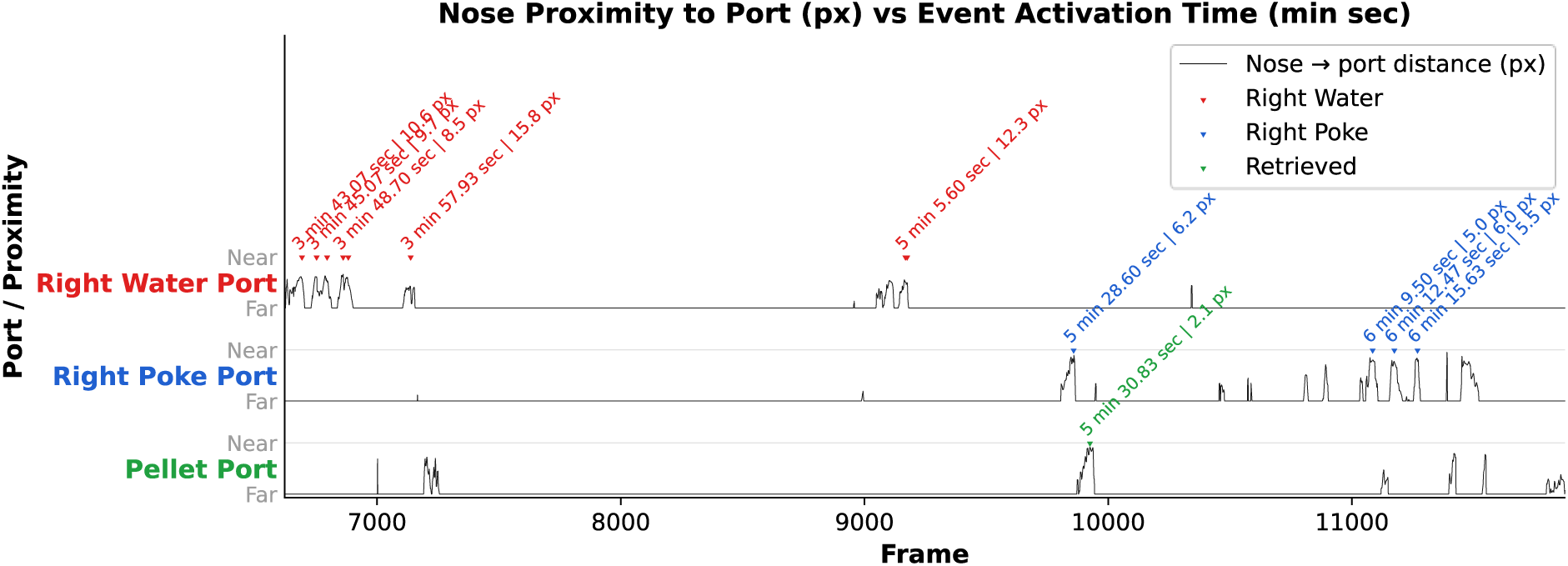
NeuroHab operant events aligned with mouse position relative to the port over 7 hours of free-moving behavior. Position data calculated from DeepLabCut (Mathis et al., 2018) analysis of camera frames output synchronously from the Mini two-photon microscope to illustrate the capacity for high-fidelity synchronization with external equipment.

**Fig. 12.**
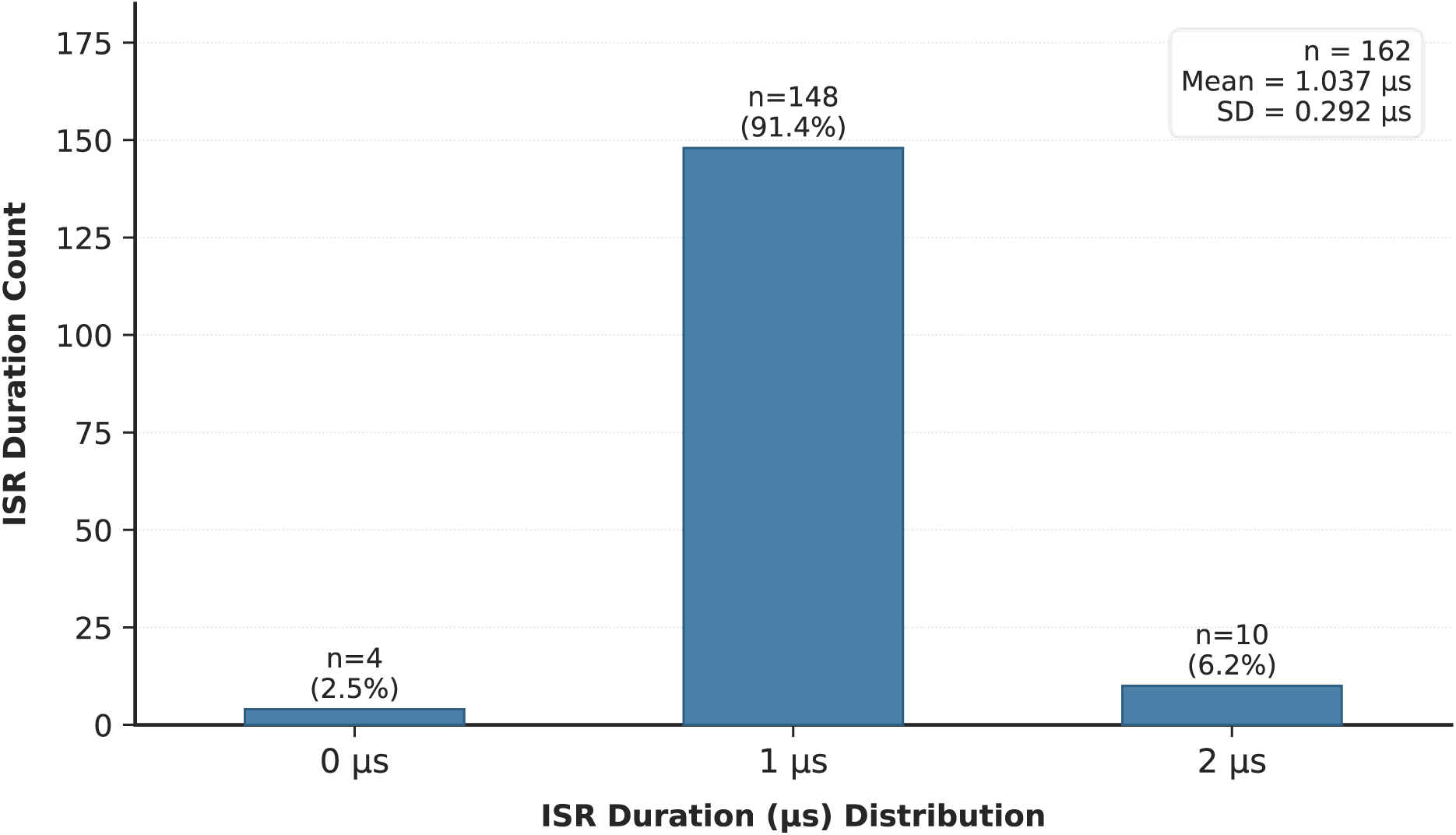
ESP32 event interrupt service routine (ISR) execution time distribution. Most ISRs complete within 1 microsecond. Illustrates the burden of event recording latency stems primarily from response latency.

**Fig. 13.**
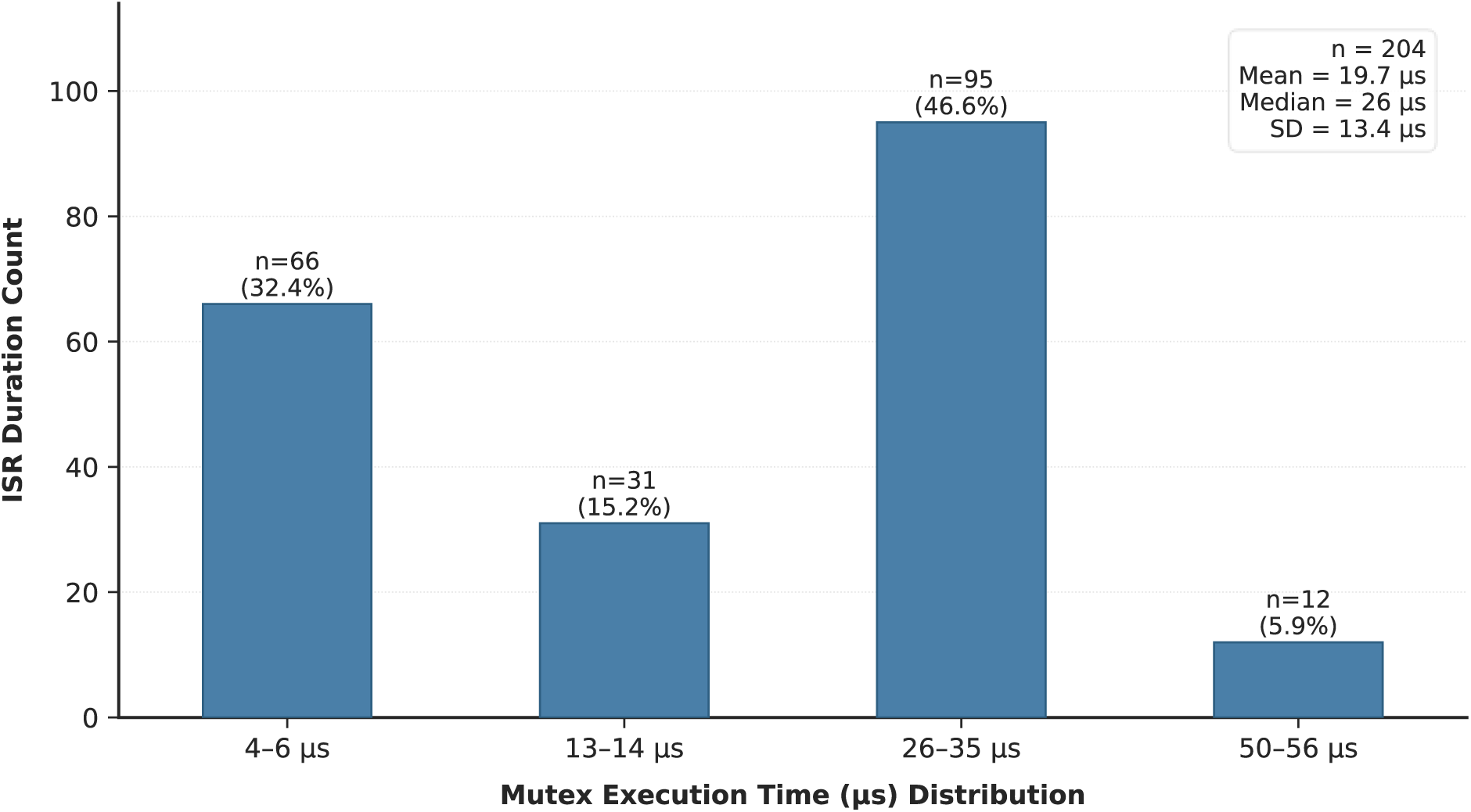
ESP32 mutex and buffer switching execution time distribution. Higher times come from buffer switching requirements on incoming pulses. Lower execution times result from non-buffer switching periods. As with figure 12, illustrates ESP32 side logging accounts for a small percentage of total behavior to log time.

**Fig. 14.**
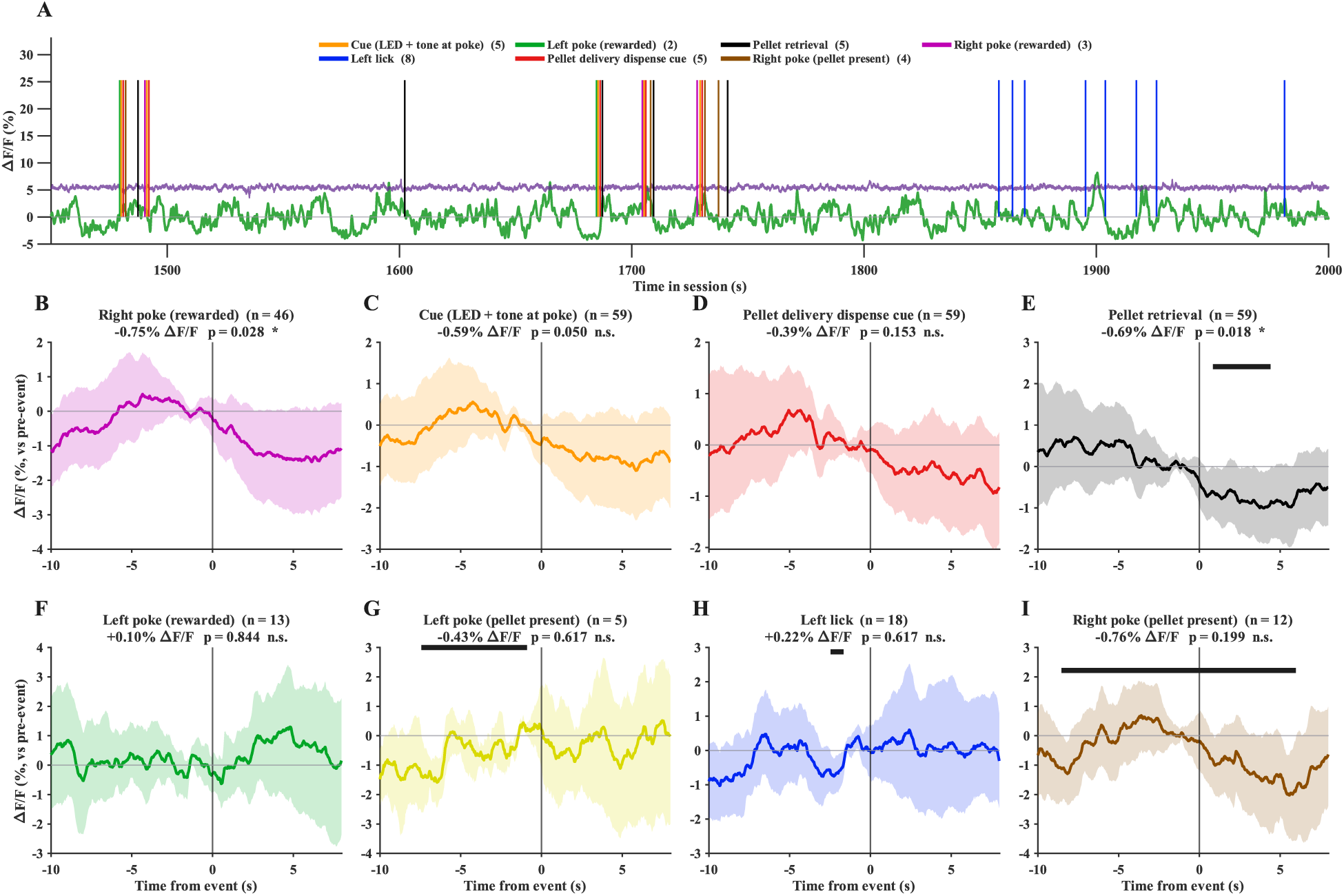
Mouse infralimbic cortex population activity aligned with NeuroHab TTL events utilizing shared BNC channel reference synchronization via NeuroHab logging architecture. **A.** Representative population dynamics (green) with isosbestic channel (purple) aligned with NeuroHab events at <1ms resolution. The isosbestic was used to detect movement artifact during the free-moving calcium imaging experiment. **B – I.** Representative peri-event traces showing event average population ΔF/F activity aligned to event onset (time = 0 s; n = events). Solid line indicates the mean; shaded region indicates ±SD.

In this paper, we describe the NeuroHab’s modular architecture and integrated control system, characterize its temporal precision and synchronization capabilities, and demonstrate its utility in combined behavioral-neural experiments. We provide detailed documentation of hardware components, software implementation, and validation procedures, to encourage other laboratories to adopt and adapt this platform for their own research needs.

## 2. Results

**Table 1.** Summary of key NeuroHab performance metrics.

| Metric | Value / Range |
| --- | --- |
| Behavior-to-log latency | <1 ms (typical 56–728 μs) |
| Max behavioral event logging rate | 16.67 Hz (single-pulse events); ~10 Hz at 39-pulse encoding |
| Sustained BNC input throughput | up to ~247 Hz single-channel (5 s window) |
| Recommended safe BNC rate (sustained, multi-channel) | <~ 13.2 Hz per channel |
| Recommended safe BNC rate (sustained, single-channel) | 47.5 Hz single channel |
| Lick detection accuracy | 96.83–100% across validation sessions |
| Liquid dispensing error | as low as 1.04% in-range; ≤2.95% across calibrated ranges |
| Approximate system cost | ~\$1,400 (all modules) |
| Deployment uptime | >50 sessions over ~6 months with no critical failures |

### 2.1 Operant Capabilities

The NeuroHab supports a range of operant conditioning paradigms through its integrated modular components (modular components, hereafter “Mods”). Food reward is delivered via the FED3 (Matikainen-Ankney et al., 2021; Nguyen et al., 2016) pellet dispenser which provides two nose-poke ports and a pellet retrieval port for operant food training. Fluid delivery is managed by up to three Lickport modules, each capable of recording activation events and dispensing calibrated liquid volumes upon contact or per programmed schedule. Auditory and visual conditioned stimuli are delivered via up to three conditioned stimulus (CS) modules, each providing an active buzzer and RGB LED. All Mod activations and stimulus deliveries are automatically recorded by the Core without additional researcher intervention. Operant schedules are programmed in C++ via the NeuroHab Arduino library, which exposes methods for Mod control and event recording without requiring modification of the underlying logging architecture. This enables the implementation of diverse paradigms ranging from simple continuous reinforcement to timed feeding, random progressive ratio, and variable interval schedules. Sample head-fixed licking trial data is available in figures 6a, 6b, and 6c, where licking rates of up to 7 Hz were recorded under a free water delivery paradigm. Mods have also been successfully integrated to replace static water and pellet delivery devices for long-term home-cage recording, with events across a continuous 16-hour session illustrated in figures 9a and 9b.

### 2.2 Temporal Precision and Event Logging

The NeuroHab Core achieves a behavior-to-log latency of less than 1 ms across all native Mods, with typical latencies ranging from 56 μs to 728 μs under normal operating conditions calculated based on response latencies and instruction execution times (Sections 3.5.1, 3.9.10 and Figures 2, 8, 12, 13). Lickport capacitive sensors exhibit mean response latencies of 223.48 μs (SD = 5.49 μs, n = 200), 424.96 μs (SD = 8.84 μs, n = 200), and 615.88 μs (SD = 9.90 μs, n = 200) with 1, 2, and 3 Lickport sensors active (Figure 2). Following NeuroHab integration, FED3 (Matikainen-Ankney et al., 2021; Nguyen et al., 2016) IR sensor response latency does not exceed 56 μs, with a mean of 28.96 μs (SD = 14.13 μs, n = 200) across 100 samples per port (Figure 8). Conditioned Stimulus modules exhibit negligible response latency as activation-only devices. Specific latency calculations are available in sections 3.5.1 and 3.9.10.

### 2.3 Integration with Neural Recording

All behavioral events logged by the NeuroHab are forwarded in real time via BNC port 1 to external data acquisition systems (covered in-depth in sections 3.4.1 and 3.5.1). This enables direct temporal alignment of operant behavioral events with neural recording including two-photon calcium imaging and electrophysiology. Behavioral timestamps are logged with <1 ms latency, and alignment with external acquisition hardware (e.g. the M2P) is accurate to within ±1 ms, bounded by the external device’s timestamp resolution (Figures 10a-10d, 14). This places behavioral events within the timescale of individual calcium transients and enables reliable inference of stimulus-response relationships at the level of single cells (Buzsáki, 2004). We emphasize that the results reported here constitute a technical validation of timing and synchronization performance. While the platform is designed to support cell-level stimulus– response analyses, detailed peri-event neural response profiles will be presented in future, dedicated studies.

The NeuroHab is compatible with Mini two-photon microscopy for simultaneous recording of prefrontal cortex activity during operant behavior. As a proof-of-concept, Figure 11 illustrates synchronization of the NeuroHab with the Mini two-photon. TTL pulses delivered through BNC port 1 served as behavioral timestamps, enabling precision post-hoc alignment of licking events, water delivery, conditioned stimuli, and nose poke events with two-photon camera frame data.

### 2.4 BNC Recording Frequency Validation

Temporal alignment across devices for precision synchronization of operant, recording, or managerial events is necessary for post hoc analysis of experimental data. NeuroHab BNC recording input enables internal synchronization with external devices for real-time alignment of events.

Single-channel BNC input recording supports sustained event captures up to 247.6 Hz, 62.4 Hz, and 47.5 Hz across 5, 60, and 120 second recording windows, respectively (n = 3 trials per condition). Dual-channel recording supports speeds up to 62.4 Hz, 16.4 Hz, and 14.1 Hz across the same windows (n = 3 trials per condition). Triple-channel recording achieves up to 27.8 Hz, 14.1 Hz, and 13.2 Hz across 5, 60, and 120 second windows as well (n = 3 trials per condition).

These represent the maximum frequencies at which all sent pulses were received across three consecutive trials for each reported Hz, channels active, and recording window (Figures 4a, 4b, 5a, and 5b). (Note: BNC rates represent discrete TTL event-pulse detection, not continuous analog waveform sampling; the NeuroHab BNC system is not designed for kilohertz-rate electrophysiology recording.)

Under staggered pulse conditions, in which inputs are offset by 50 μs, maximum capture frequencies increase substantially across dual and triple channel configurations. Dual-channel recording achieves up to 162.8 Hz, 89.7 Hz, and 61.9 Hz across 5, 60, and 120 second windows respectively (n = 3 trials per condition) and triple-channel recording achieves up to 161.4 Hz, 61.7 Hz, and 47.2 Hz across the same windows (n = 3 trials per condition). These results indicate that synchronous pulse delivery represents a lower bound on system performance, and that practical multi-channel recording scenarios, in which simultaneous exact-coincidence events are rare, support higher sustained throughput. It is worth noting that dual-channel recording frequencies outpacing single channel recording at the 60s and 120s window is due to the 50 μs delay used to stagger pulses, resulting in a longer inter-event interval.

### 2.5 Lickport Accuracy

Lickports were developed as a wireless (i.e., untethered, the mouse has no electrical connection through the animal) alternative to conventional wired lick detection systems, offering a form factor analogous to a standard home cage water bottle spout to promote naturalistic licking behavior. Detection accuracy and dispensing precision were validated across multiple independent trials; full protocol details are provided in Methods Section 3.8.1.

Lickport detection accuracy was assessed across three independent validation sessions achieving accuracies of 100%, 100%, and 96.83% respectively (Table 2). The single session falling below 100% accuracy (96.83%; n = 63 activations) was attributable to camera frame-rate limitations during video validation rather than confirmed system error. In a free-moving trial, licking bout frequencies averaged 0.58 Hz for bouts under 10 seconds, 2.55 Hz for bouts under 1 second, and 4.27 Hz for bouts under 0.5 seconds, with a maximum instantaneous frequency of 14.49 Hz (69 ms inter-lick interval). Licking frequencies in the head-fixed trial were consistent with the free-moving trial, averaging 0.51 Hz, 2.96 Hz, and 3.46 Hz for the same bout duration ranges, with a maximum instantaneous frequency of 7.06 Hz. The lower maximum frequency observed in the head-fixed condition is attributed to the postural constraints imposed by head fixation relative to free movement. These results demonstrate that the Lickport system provides reliable, high-accuracy lick detection comparable to wired approaches while eliminating the constraints associated with physical tethering. In the free-moving condition, 100% accuracy indicates that all 105 logged lick events corresponded to observable licking behavior in the video recordings. Because limited camera resolution and frame rate can cause some individual lick contacts to be missed in the top-down view, the total number of actual licks may be underestimated, making the reported accuracy a conservative estimate.

Lickports are additionally capable of dispensing precise volumes of liquid via solenoid valve actuation, with dispensing error as low as 1.04% when the system is calibrated to the active reservoir range. Dispensing error remains below 2.95% across calibrated ranges and increases when dispensing occurs outside the calibrated reservoir range, with errors reaching up to 9.37% in uncalibrated conditions due to reservoir-height-dependent pressure variation. Taken together, these findings support the use of Lickports as a high-accuracy, wireless solution for both lick detection and precise liquid delivery in rodent behavioral paradigms. Users are advised to calibrate to their active reservoir range and to periodically verify dispensing accuracy with mass-based checks, particularly when using volumes outside the validated range.

### 2.6 System Deployment and Neural Synchronization Validation

The NeuroHab has been deployed across more than 50 behavior trials as the primary data collection device. In over 20 experiments, the system was operated in tandem with the Mini two-photon microscope (M2P) to correlate operant behavior with prefrontal cortex activity, with behavioral event timestamps synchronized to M2P imaging frames via BNC port 1.

Random progressive ratio, free feeding, 2×2, and 5×5 port switching paradigms, among others, have been performed with mice in the NeuroHab system. For standalone experiments, all data is recorded on the NeuroHab in.csv format with sub-millisecond timestamps for later analysis (Figures 9a, 9b). This.csv contains the timestamps and counts for the session from which required data (like retrieval time) may be extrapolated. When paradigms are performed with M2P imaging, the.csv data is also exported simultaneously to the M2P system for recording via BNC port 1. Because the NeuroHab can use the M2P as a shared reference, we can align the NeuroHab timestamps with calcium imaging (or other methods, see TTL alignment with fiber-photometry prefrontal population activity in Figure 14) post hoc to see precisely when events occurred relative to camera frame data (Figure 11). This alignment framework supports quantitative analysis of relationships between neural activity and operant behavior at the level of individual events and cells (Qian et al., 2026).

Synchronization fidelity between the NeuroHab and M2P systems was assessed by comparing the interval between consecutively received pulse train timestamps at each device. Because both devices receive pulses from the same physical wire, any measured difference in inter-pulse interval reflects timing artifacts introduced by each device’s internal clock rather than a genuine difference in event timing. The distribution of interval differences (Figure 10a) was centered at a mean of −22 µs with a standard deviation of approximately 482 µs, indicating negligible systematic offset between devices. The spread of the distribution is bounded by ±1000 µs and reflects the M2P’s 1 ms timestamp resolution: because the M2P records arrival times to the nearest millisecond, the difference between any two NeuroHab and M2P interval measurements carries up to ±1000 µs of quantization noise regardless of true synchronization quality. The slight left-tail asymmetry in the distribution (extending to approximately −8900 µs) is attributable to free-running clock drift, whereby the relative rate difference between the NeuroHab and M2P clocks accumulates over time between events, with longer inter-event intervals producing proportionally larger discrepancies. This drift is supported by the scatter plots in figures 10b– 10d: when intervals below 1 second are excluded, removing the area where quantization noise dominates, the log-linear relationship between interval length and absolute timing difference strengthens markedly (R² rising from 0.074 to 0.155), and when intervals below 1 minute are excluded, the relationship strengthens further to R²=0.651 (Spearman ρ=0.415, p<0.001). The apparent discrepancy between the steadily rising R² and the non-monotonic pattern in Spearman ρ (0.293 in Figure 10b, falling to 0.123 in Figure 10c, then rising to 0.415 in Figure 10d) reflects the disproportionate weight of quantization noise at short intervals, not a fit driven by outliers. Because the majority of inter-event intervals fall below 1000 ms, and because the fixed 1 ms quantization step is comparatively large relative to both interval length and interval difference in this range, this densely populated short-interval region in Figure 10b artificially strengthens the monotonic relationship (ρ=0.293). When intervals below 1 second are excluded in Figure 10c, this quantization-dominated mass, which had been propping up the correlation, is removed, and the remaining long-interval observations, now more susceptible to noise from individual outliers, produce a temporarily weaker relationship (ρ=0.123). As the threshold is extended further, to intervals greater than 1 minute in Figure 10d, the fixed 1 ms quantization step becomes small relative to both the interval length and the accumulated drift, so its distorting influence recedes and the true drift relationship reemerges more clearly (ρ=0.415).

This pattern would be expected to continue at longer thresholds still (e.g., 5, 10, or 30 minutes), as the relative contribution of quantization noise continues to shrink. As each interval difference is measured independently from its own start time, the clock drift does not accumulate across trials. Taken together, these results confirm that the NeuroHab and M2P systems are effectively synchronized within the limits imposed by M2P’s millisecond recording resolution, and that observed inter-device discrepancies are fully explained by quantization noise at short intervals and clock drift at long ones.

Over approximately six months of routine deployment, more than 50 behavioral sessions were run without critical system-level failures; no loss of event logs or complete timestamp corruption was observed. Observed issues were limited to non-critical hardware problems: temporary solenoid valve sticking (resolved by priming the line and removing air bubbles) and occasional loose BNC connectors (mitigated by securing cables and adding strain relief). Systematic lifetime characterization, for example, solenoid actuation cycle counts and PCB longevity, has not yet been fully quantified; users planning chronic, high-throughput studies are therefore advised to implement routine hardware inspection and maintenance schedules.

## 3. Materials and Methods

### Ethical Approval

All animal experiments described in this study were approved by the Institutional Animal Care and Use Committee (IACUC) of the University of Wyoming (protocol number #2022-0127) and were conducted in accordance with national guidelines for laboratory animal welfare, including the NIH Guide for the Care and Use of Laboratory Animals.

### 3.1 System Architecture Overview

NeuroHab architecture consists of two physical components: the behavioral enclosure (hereafter “Home”, Figure 1 mouse enclosure) and the central control unit (hereafter “Core”, Figure 1 control box with BNC ports and LCD Display). The Home is where the mouse is housed and contains the NeuroHab modular components (“Mods”, Figure 1 see Mods in right legend) that the mouse interacts with. The Core is the hardware integration system which controls Home behavior. The Core consists of a dual microcontroller architecture, Arduino MEGA 2560 (Arduino, 2023a) and ESP32 DevkitC (Espressif Systems, 2024b). The Arduino is responsible for executing Home behavioral paradigms and the ESP32 for logging events to the onboard micro-SD card. One-way device communication, Arduino (control) to ESP32 (logging), is accomplished through TTL pulses. The two devices are connected via custom PCB (printed circuit board) which allows for the integration of operant Mods. System source and documentation are available in the accompanying repository (Sun Lab, 2026).

### 3.2 NeuroHab Home

Mouse interactions occur in the Home which consists of a modular 3D printed base with walls and floors made of 6mm acrylic glass. The 3D printed base contains mounts for the installation of cameras or lights on the attached swing arms. The base also contains a dual reservoir holder for 60ml syringe barrels for drug/liquid storage and delivery. All floors and walls are removable for easy cleaning and modification. The Home is a blank slate for Mod installation and modification to suit the experimental design. Further, Mods may be installed within the home cages of mice and proprietary setups to utilize the Core separately from the Home.

### 3.3 NeuroHab Core, Open Source and Modifiability

The NeuroHab is designed as a highly modifiable, open-source platform tailored to meet the diverse needs of neuroscience and behavioral research laboratories. By providing robust event-logging integrated with an extensible operant conditioning toolkit, it supports a wide array of behavioral paradigms and experimental configurations. The system’s architecture enables seamless control of external hardware components beyond the core modules described here. As a result, researchers can readily adapt the NeuroHab, or implement entirely novel experiments, with minimal additional development effort: the underlying data acquisition, timestamping, and control logic are already in place, allowing focus to remain on scientific innovation rather than low-level systems engineering.

Modular components, or Mods, serve as the primary interactive elements within the Home environment, delivering standard reinforcers and stimuli such as food, water, drugs, light, and sound (detailed in 3.8 NeuroHab Mods). Recognizing that many labs require specialized or custom hardware (Akam et al., 2022; Kapanaiah et al., 2021), the Core was explicitly engineered for straightforward external Mod integration. The onboard Arduino provides approximately 30 available digital and PWM pins, offering ample flexibility for custom additions. Users can incorporate their own Mods while leveraging the existing logging system with little effort. For detailed implementation guidance, including pin assignments, code examples, and best practices, refer to our tutorials on external Mod integration (Sun Lab, 2026).

The NeuroHab Core is responsible for the control flow of Home behavior and the logging and external synchronization of behavior events. Core architecture revolves around a dual microcontroller setup. The onboard PCB connects the Arduino MEGA 2560, ESP32 DevkitC, internal components, and Mods reproducibly. The dual microcontroller setup separates event control flow and logging at the hardware level to improve precision recording capabilities and reduce design complexity. The standardization of these modules reduces the technical complexity of rig development for researchers by abstracting necessary components such as logging, control flow, and paradigm design without reducing usability.

### 3.4 Core, Arduino Mega 2560

The Core Arduino is responsible for directing control flow of experimentation, integration of Mods, and Mods’ TTL communication with the logging system. The Arduino implements a custom NeuroHab library written in C++. This library contains methods for interacting with NeuroHab Mods and recording events without directly modifying source code. Utilizing this structure enables behavior paradigm creation without reprogramming logging or control flow schema, improving confidence in experiment execution and reducing time spent on redevelopment and validation.

#### 3.4.1 Arduino TTL Communication

The Arduino generates TTL pulses to communicate with the ESP32-based logging system for event recording. The number and types of events are modifiable by the researcher within system limitations.

TTL communication connects ESP32 pins to ground to generate falling edges a specific number of times (n pulses), within a timeout window, and with a fixed inter-pulse delay (500 μs). The number of pulses, n, encodes specific events sent to the ESP32 for equivalent decoding and logging. This encoding system improves precision, decreases pin requirements, reduces interrupt complexity, and improves system robustness. The current event encoding for TTL pulse trains is as follows (Table 2):

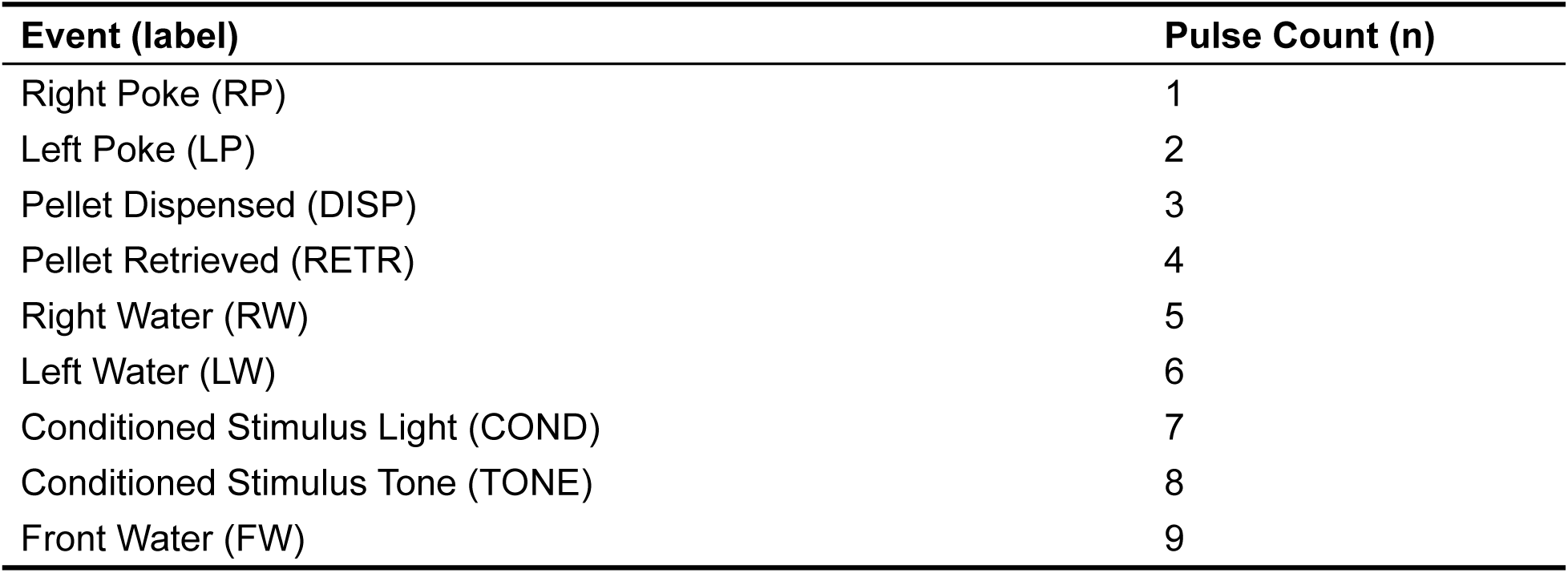
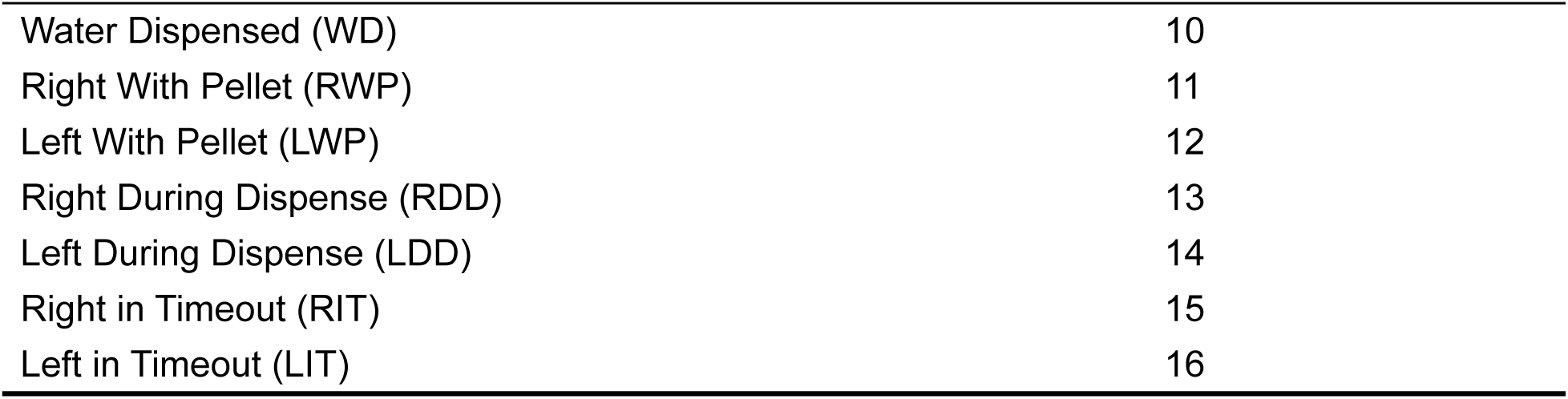

| Event (label) | Pulse Count (n) |
| --- | --- |
| Right Poke (RP) | 1 |
| Left Poke (LP) | 2 |
| Pellet Dispensed (DISP) | 3 |
| Pellet Retrieved (RETR) | 4 |
| Right Water (RW) | 5 |
| Left Water (LW) | 6 |
| Conditioned Stimulus Light (COND) | 7 |
| Conditioned Stimulus Tone (TONE) | 8 |
| Front Water (FW) | 9 |
| Water Dispensed (WD) | 10 |
| Right With Pellet (RWP) | 11 |
| Left With Pellet (LWP) | 12 |
| Right During Dispense (RDD) | 13 |
| Left During Dispense (LDD) | 14 |
| Right in Timeout (RIT) | 15 |
| Left in Timeout (LIT) | 16 |

For clarity, when the mouse licks the right water port for water delivery the following will occur. The Arduino cycles paradigm code until NH.RW_lick() returns true. NH.RW_lick() will connect communication pins 25 and 26 to ground 5 times in approximately 5 ms to signal BNC out and ESP32 communication. This is followed by a pulse separation delay further outlined in section 3.9.3. The ESP32 will interrupt its current behavior on the first pulse and save the time since initialization and the event which occurred. This occurs in under a millisecond. More on how the ESP32 logs events shortly (section 3.5).

#### 3.4.2 Extending the System with Custom Events and Modules

The TTL encoding scheme is designed to be extended by end users who wish to add new behavioral events or integrate additional external modules. To add a new event, the user (i) selects an unused pulse count within the 1–39 range, (ii) adds the corresponding mapping to the Arduino-side encoding definition in NeuroHab_Control.h, (iii) duplicates the new values to the ESP32-side decoder mapping in the logging firmware in NeuroHab_ESP_Logging.ino so that the new pulse train is interpreted as the corresponding event code, and (iv) runs a short test schedule to verify correct encoding, decoding, and timing of the new event. Concrete code templates and example modules are provided in the project’s GitHub repository (Sun Lab, 2026) under Tutorials/Event Encoding as starting points for customization. This workflow allows the multi-module architecture to be expanded without modifying the core logging pipeline.

#### 3.4.3 Arduino C++ Control Scripts

Control of NeuroHab behavior and paradigm execution is standard C++ Arduino logic. The NeuroHab is initialized in the setup() method and logic is run repeatedly during loop(). This looped logic controls the behavior paradigm created by the researcher. For example, a free licking paradigm (Figures 6a, 6b, 6c) delivers a set amount of liquid to the mouse every Lickport activation. Conversely, a random progressive ratio licking paradigm requires multiple and variable Lickport activations to receive the same quantity of water. The control scripts are capable of all component control schema for the development of novel operant paradigms in accordance with researcher requirements.

### 3.5 Core, ESP32 DevkitC

The ESP32 is responsible for the precision logging of Arduino events and externally connected pulses. The ESP32 has single and multi-channel logging capabilities for external devices and up to 16.67 Hz logging capabilities for Lickports and other NeuroHab Mods under default logging configurations. The ESP32 records approximately 16 operant events per second at <1 ms latency. BNC input captures TTL event pulses (discrete event detection) at up to 247 Hz on a single channel. This 247 Hz rate refers to event-pulse detection only, not continuous analog waveform sampling; the NeuroHab BNC system is not designed for kilohertz-rate electrophysiology waveform recording. The NeuroHab utilizes 1 to 3 channel BNC input recording with high interval capabilities (76.63 Hz to 247 Hz) for single channel logging and low interval capabilities (13.2 Hz) for simultaneous triple channel logging under longer pulse trains, worst case (Figures 4a, 4b, 5a, 5b).

The primary BNC port on the Core forwards all ESP32 received communication, with no additional latency, for synchronization with external data collection modes (Figures 10a-10d). Logging is integrated in NeuroHab-ESP_Logging.ino written in C++, whose architecture and design decisions are discussed in the following section.

#### 3.5.1 Architecture and Design of NeuroHab-ESP Logging for Precision Logging

Three software principles have been integrated within the logging system to achieve the precision required for neural data integration. These are the batching and switching of incoming behavior events for later writing (Tanenbaum and Bos, 2014), utilizing pin interrupts on ESP32 hardware (Espressif Systems, 2024a), and ESP32 hardware mutex locks (Espressif Systems, 2024a; Tanenbaum and Bos, 2014). To briefly overview, incoming events are stored in ESP32 memory on three pairs of 2D arrays which hold the necessary data, until the processor is free to write the events to disk.

The purpose of batching and switching of incoming data is to provide a buffer between memory and disk (Tanenbaum and Bos, 2014). As it takes far longer to write to disk than to memory, it is impossible to record fast events directly to disk without losing data. The solution is to save incoming events to memory while other events are written to disk. These arrays account for the number of pulses received, the microsecond timestamp of the first pulse received, and the microsecond timestamp of the most recent pulse received. Each pair of arrays consists of an A and B array. While A buffers are being written to disk, B becomes the active buffer to hold incoming events. Once B becomes full, A becomes active while B is written to disk. Here lies the limit on the maximum Hz at which we can record via BNC. We may record at higher frequencies by increasing the size of our buffers, but the maximum lies at the hardware memory limit.

Pin interrupts are used to signal devices to execute an Interrupt Service Routine (ISR) (Espressif Systems, 2024a; Tanenbaum and Bos, 2014). We implement ISRs to fill the buffers with the necessary incoming data for event decoding. Our ISRs utilize timeouts to separate incoming logic for the decoding of behavior information. The ISR also checks validation details like pulse length to prevent erroneous, noisy data. Once the ISR completes, there is no additional latency between behavior and recording timestamps as the microsecond timestamp is saved for later writing. Once a Mod’s sensor detects an event, it takes only a few microseconds to send the first ground pulse from the Arduino to the ESP32. The ESP32 takes a few more microseconds to recognize the pin interrupt and jump to the ISR, whose time complexity is O(1) and runtime is less than 3 μs (Figure 12). While the first timestamp is nearly instantaneous, the sending and decoding of specific events takes about one additional millisecond per pulse sent, plus a separation delay. This leads to the 16.67 Hz behavior event recording limitation which we will discuss further in section 3.9.3.

The process of writing buffered data to disk is governed by classic challenges in asynchronous and multithreaded programming (Tanenbaum and Bos, 2014). To address these, we implement buffer switching within a critical section protected by a mutex lock. Specifically, the commands portENTER_CRITICAL(&mux) and portEXIT_CRITICAL(&mux) (Espressif Systems, 2024a) obtain and release our mutex, delaying any ISRs and shielding the buffers from interference while the enclosed code executes. Within this critical section, we employ memcpy and memset to copy buffer data to local arrays and reset the buffers to empty states (Figure 7). On each loop iteration, we acquire the mutex locks and check whether a buffer switch is required; if so, we perform the switch. Any incoming data is stored in local arrays and written to disk once the CPU becomes available.

The following outlines representative scenarios and their associated timing and event-handling procedures. For simplicity, we denote behavioral events as X, Y, and Z. We make the following non-impactful assumptions to facilitate readability: the ESP32 begins with buffer A active and executes from the start of its loop; the ESP32 loops repeatedly while awaiting events (with negligible loop time, as ISRs remain active); and event X is transmitted first from the Arduino to the ESP32 for logging.

The external ground pulse connects the ESP32’s pin 4 or 5 to ground, triggering an ISR. The transition to the ISR and its initial timestamp logic require less than 3 μs (Figure 12). Additional event data describing X is then recorded in buffer A. The ISR concludes, and execution resumes where the loop left off. The loop persists until the buffer fills or sufficient time elapses. At this point, the critical section is entered, the active buffer is copied to local arrays, the critical section exits, and logging proceeds.

If event Y occurs during buffer switching, it remains pending until the buffer switch completes (typically around 30 μs, and always less than 56 μs) (Figure 13). Event Y is then written to buffer B, after which event X is committed to disk. If event Z arises while Y is being written to disk, the ISR activates, Z is recorded in the active buffer, and the ISR returns to complete Y’s disk write without data loss, thanks to timestamps preserved in local arrays. In the worst-case scenario, where Y coincides with buffer switching, the delay to Y’s timestamp is less than 56 μs (Figure 13). In practice, Y must follow X by less than 20 ms (60 ms batch-write timeout − 40 ms BNC event timeout) for this loss to accrue.

Latency calculations for ESP32 hardware were evaluated from 148 samples of operant ISR latency and 204 samples of critical section execution time as these are the only points of execution which could introduce latency to incoming events. Our measurements in Figure 12 indicate mean execution latencies for the ISR responsible for recording initial timestamps at 1 μs for 91.4% of events and 2 μs as the maximum. The latency introduced by buffer switching described in the previous paragraphs is shown in Figure 13, with critical section execution times evaluated between 4 μs and 56 μs. The latency for the system is calculated as the response latency of the sensor added to the logging latency of the ESP32. For CS modules with no response latency, the system latency is approximately 1 μs if it is the first recorded event and less than 56 μs if the event is pended due to buffer switching. For triple Lickport response latencies, the system latency in the worst case is approximately 728 μs (672 μs outlier from 3 Sensors Figure 2 + 56 μs worst case buffer switching latency Figure 13).

### 3.6 Core, BNC I/O

The NeuroHab Core integrates five BNC I/O ports for synchronization and communication with external hardware. BNC port 1 serves as the primary output, forwarding all TTL behavioral event pulses generated by the ESP32 for direct, real-time, integration with neural recording systems such as two-photon imaging and electrophysiology setups. BNC port 2 provides an unformatted TTL output channel, allowing researchers to interact with external devices’ BNC input independent of NeuroHab event pulses. BNC ports 3 through 5 function as input recording channels, enabling the NeuroHab to receive and timestamp TTL pulses from external devices for synchronization with other data collection modalities. Input recording leverages the ESP32 buffering architecture described in Section 3.5.1 to achieve sustained capture rates of up to 247 Hz on a single channel, with simultaneous triple-channel recording supported at rates up to 13.2 Hz under worst-case synchronous pulse conditions (Figures 4a, 4b, 5a, 5b). All incoming BNC events are saved to a dedicated .csv file on the onboard micro-SD card, which shares a common reference time with all other NeuroHab behavioral and neural data events, enabling straightforward post hoc alignment across modalities.

### 3.7 Integration with Neural Recording and Synchronization Validation

The precision recording capabilities of the NeuroHab were developed specifically for integration with neural recording methods such as calcium imaging (Zong et al., 2017) and electrophysiology. All behavioral events logged by the NeuroHab are forwarded via hardware to BNC port 1, allowing direct connection with external imaging systems. This positions the NeuroHab as an intermediary, synchronizing outputs from operant devices, many of which lack native compatibility with neural acquisition hardware, with neural data, without introducing latency.

Temporal synchronization between the NeuroHab and Mini two-photon (M2P) recording systems was validated using event data collected across a single 16-hour free behavior trial. During this trial, behavioral events generated by the mouse were recorded concurrently and independently by both systems: the NeuroHab system logged event timestamps internally, while the M2P system saved the same pulse trains for later event evaluation via comparison to the event decoding table to identify corresponding event types and their onset times. Inter-event intervals were then calculated for each system independently as the elapsed time from the start of one event to the start of the immediately following event, producing two matched series of interval durations, one per device, across the full trial. Synchronization was assessed by computing the difference between NeuroHab and M2P interval durations for each matched event pair (NeuroHab interval − M2P interval), with differences converted to microseconds to resolve sub-millisecond structure (Figure 10a). The resulting difference series was analyzed for systematic offset, distributional spread, and the relationship between interval duration and absolute timing discrepancy, the latter being used to distinguish between the M2P’s fixed 1 ms quantization noise floor and the duration-dependent accumulation expected from free-running clock drift (Figures 10a-10d).

With <1 ms behavioral event logging latency and millisecond-scale synchronization to the M2P (bounded by the M2P’s 1 ms timestamp resolution; see Figure 10a), the NeuroHab enables temporally precise correlation of operant behavior with neural activity. (Buzsáki, 2004). Our system is compatible with electrophysiology setups by generating analog falling-edge pulses detectable as discrete spikes in recordings, ensuring that behavioral markers appear alongside neural traces.

The NeuroHab’s low cost and versatility reduces barriers to combining operant behavioral experiments with high-fidelity neural analysis. Researchers can move beyond traditional operant conditioning paradigms to conduct temporally synchronized multimodal studies with the integration of diverse experimental design normally unsuited for combination with neural analysis.

### 3.8 NeuroHab Mods, Design and Integration

NeuroHab modular components (Mods) serve as the primary tools through which researchers design operant schedules. The current NeuroHab implementation supports three categories of integrated Mods. The Lickport module detects animal contact and delivers calibrated fluid volumes upon activation or according to a programmed paradigm. Conditioned stimulus (CS) modules provide auditory and visual stimuli via an active buzzer and RGB LED, respectively. Food reward is administered through the FED3 pellet dispenser developed by the Kravitz Laboratory (Matikainen-Ankney et al., 2021; Nguyen et al., 2016) and modified for integration with the NeuroHab control and logging architecture. Each Core supports up to three Lickport modules, three CS modules, and one FED3 unit. Researchers may extend this framework by incorporating additional Mods or external operant components per experimental requirements.

#### 3.8.1 Mods, Design of Precision Liquid Dispensing Ports (Lickports)

The Lickport was designed to provide rodents with a drinking interface similar to the standard home cage water bottle spout, while integrating precise capacitive lick detection and solenoid-controlled liquid dispensing. Each Lickport unit combines a dispensing needle, a solenoid valve, and a capacitive touch sensor to deliver quantified volumes of liquid upon mouse contact. Lickport events and all subsequent dispensing actions are logged within the Core, and Lickport behavior is fully programmable within NeuroHab control scripts to define the timing, conditions, and volume of liquid delivery. The system supports lick event recording at rates of up to 16.67 Hz (section 3.9.3) in parallel with dispensing.

Capacitive Lick Detection: Lickport sensors detect mouse contact via changes in circuit capacitance. When the mouse contacts the dispensing needle, the overall capacitance of the circuit rises; this change is detected by the onboard Arduino controller using the CapacitiveSensor library (Badger and Stoffregen, n.d.) and subsequently activated. TTL data is then relayed to the ESP32 for logging. To reduce noise and interference across cable length, insulated and shielded cabling is used throughout the sensor circuit. A rolling buffer of recent capacitance readings is maintained continuously, from which a dynamic baseline average is calculated. A user-defined threshold determines the minimum capacitance rise required to register an activation. Activations are triggered on the rising edge of capacitance, and a subsequent activation is suppressed until a falling edge is detected, that is, until the sensor value returns within a defined tolerance of the baseline. This rising/falling edge architecture enables reliable discrimination of individual lick contacts and prevents double-counting during sustained contact events. The system self-calibrates over time as the baseline dynamically adjusts to ambient conditions, improving robustness across sessions.

Validation Protocol: Detection accuracy was validated across three independent sessions. First, 150 manual benchtop activations were performed using a human finger across four contact conditions: touch-and-release, touch-and-hold, rapid touch-and-release, and all preceding conditions on both dry and wet surfaces (Figure 3). All 150 activations were detected, yielding 100% accuracy. Second, an 8-hour free-moving trial was conducted in which Lickports served as the sole drinking source. Of 105 activations recorded on the right Lickport, all 105 were confirmed as genuine licking events via video review, yielding 100% accuracy. Third, a head-fixed licking trial was analyzed via video review of 63 discrete activations, of which 61 were confirmed as distinct contact events; the 2 unconfirmed events are attributed to camera frame-rate limitations and may represent legitimate activations occurring between frames, yielding a conservative accuracy estimate of 96.83% for this session.

Dispensing Validation: Liquid dispensing volumes were validated by comparing predicted and delivered volumes as the mass of the dispensed liquid. At a reservoir height of 685 mm above the Lickport, calibrated across a dispensing range of approximately 60 to 5 ml, mean dispensing error across three trials was 0.27% (range: −1.58% to 1.36%), with total dispensed volumes of 53.52, 53.69, and 53.29 ml. When dispensing smaller volumes (approximately 9 ml per trial), error increased modestly: trials calibrated to the 60–51 ml range yielded errors of 1.41–2.64%, while trials calibrated to the 20–11.5 ml range yielded errors of −2.51% to −2.95%. Dispensing error is primarily driven by reservoir-height-dependent line pressure, which varies across the fill range of the reservoir. When liquid is dispensed outside the calibrated reservoir range, pressure deviations cause systematic over or under-delivery, with errors reaching up to 9.37% in uncalibrated conditions. Calibration to the active dispensing range is therefore recommended to maintain dispensing accuracy within acceptable limits and to account for hydrostatic pressure changes across the reservoir fill range. When operating outside the validated volume range, users should perform a simple mass-based check (weighing the dispensed liquid) to confirm accuracy before data collection.

#### 3.8.2 Mods, Food Delivery via FED3 (Food Experimentation Device 3, Kravitz Lab)

A pellet delivery method has been established by Alexxai V. Kravitz at the Kravitz Lab at the Washington University School of Medicine in the form of the FED3 (Matikainen-Ankney et al., 2021; Nguyen et al., 2016). The FED3 is a standalone pellet dispensing device for food delivery or operant training. The FED3 utilizes IR nose poke sensors to allow mice to poke for pellet retrieval. Alone, the FED3 is sufficient for food delivery and event recording, but it is not precise enough for our neural integration and diverse behavioral experimentation. Modified source code enables the FED3 to serve as an operant pellet delivery method capable of utilizing the NeuroHab’s precision recording and external system integration capabilities.

The FED3 utilizes an Aux output for TTL communication which we connect directly to the ESP32’s pins 5 and GND. The modified FED3 source then communicates using the NeuroHab encoding system to pass FED3 events to the NeuroHab logging system by producing active-low TTL signals. The modified FED3 source produces TTL behavior events before FED3 logging takes place to reduce recording latency. In this way the actual latency of FED3 event recording is in line with NeuroHab latencies of <1 ms. Further, all FED3 events are sent to the external BNC communication port with all other Mod communication, providing an operant feeding system capable of integration with neural data collection methods.

#### 3.8.3 Mods, Conditioned Stimulus Modules

NeuroHab conditioned stimulus (CS) delivery may be administered from 3 dedicated modules and the FED3. Each NeuroHab CS module consists of an RGB LED for visual stimulus and active buzzer for auditory stimulus. Both FED3 and CS modules are programmable in C++ via their respective scripts. Conditioned Stimulus modules are connected via 5-pin cables to Arduino Core PWM pins. Currently, only LED and TONE are recorded for all conditioned stimulus modules; however, TTL modification for specific module activation is possible or easily inferable based on paradigm design.

### 3.9 System Limitations and Considerations for Operant Schedule Combination with Neural Recording

The NeuroHab’s low-latency performance, scheduling flexibility, and cost requirements were achieved through deliberate hardware and software design choices, each introducing associated limitations that researchers should consider when modifying the system. The following are key considerations when making large modifications to avoid tradeoffs in latency and performance.

#### 3.9.1 Active-Low (Pulse Drain) TTL Communication

The NeuroHab employs active-low (pulse drain) TTL communication, in which signals are transmitted by connecting the relevant pins to ground to generate falling edges. This was selected for its inherent noise resistance, as default-high lines are less susceptible to erroneous triggering, and for compatibility with the Mini two-photon microscope, which expects the same signaling standard. FED3 source code was modified accordingly to output active-low signals through its auxiliary port for integration with the NeuroHab Core.

#### 3.9.2 TTL Communication Limitations and Advantages vs Serial Communication

The NeuroHab implements TTL pulsing as the primary method of communication between microcontrollers over the alternative of serial communication. The ease of implementation of TTL better suits the open-source, low-cost model for the NeuroHab and offers significant performance and latency advantages at the cost of variable data transfer.

TTL communication encodes data by transmitting a defined number (n) of rising or falling edges along a connection, where the pulse count corresponds to a specific event. Because the first pulse in a train is detected and timestamped within microseconds, in parallel with the transmission of subsequent pulses, this approach achieves extremely low latency between event occurrence and log registration. However, the encoding scheme is constrained by the nature of the data it can transmit. Only discrete, pre-defined events may be communicated, as both the Arduino and ESP32 must share a hardcoded map of pulse counts and their corresponding event labels. Continuous or variable-valued data cannot be transmitted directly; such values can only be approximated by binning them into a fixed number of hardcoded categories, each assigned a distinct pulse train.

Conversely, this is the strength of the serial logging approach, as any value may be serialized and sent over the connection. After opening a serial connection, bits may be sent and received with transmission times dependent on baud rate and message length; for example, at 115200 baud, a 4-byte message requires approximately 347 μs for transmission, not including software overhead (Arduino, 2023b). With this approach, arbitrary messages may be sent to the logger for handling. The latency introduced by opening and closing serial connections and the transmission of various data types does not make serial logging suitable for integration with neural recording, however. While theoretically possible to timestamp incoming serial “start” bits for 4-byte data at sub-1 millisecond utilizing high baud rates, the practicality of implementing and managing the complex interrupt service routines is neither suitable for a general research audience nor easily extensible for more complex data types such as strings.

Several approaches are available to work within this constraint. Known values may be hardcoded explicitly and tracked via counters in the logging script; the Lickport modules, for instance, calculate total fluid delivery volume by multiplying a calibrated dispensed quantity by the number of activations. Custom value bins may also be defined by assigning pulse train counts to researcher-specified categories within the existing TTL encoding-decoding framework. For applications requiring variable data transmission, the intrepid user may connect the RX and TX pins of the onboard microcontrollers (Arduino, 2023a; Espressif Systems, 2024b) to enable serial logging alongside TTL communication, with TTL used to timestamp the onset of each serial transmission and the subsequent data offset accordingly, though this approach incurs latency and resolution costs that must be accepted. For neural integration experiments, our recommended approach is to timestamp all behavioral events via the NeuroHab in parallel with external data acquisition and align any variable data post hoc during analysis.

#### 3.9.3 TTL Limits Continued, Pulse Train Limits on Recording Frequencies and Synchronous Mod Activations

By default, each NeuroHab TTL pulse occupies 1 ms, consisting of 500 μs low followed by 500 μs high. Pulse train length, therefore, scales directly with the encoded event: a right poke requires a single 1 ms pulse, while a conditioned stimulus tone requires 8 ms across 8 pulses. The TTL communication timeout is hardcoded at 40 ms, placing a practical upper limit of 39 encodable events, the 40th pulse may be unreliable due to accumulated latency and signal loss at the boundary of the timeout window. A mandatory transmission separation delay (PULSE_SEPAR_DELAY_MS) of 60 ms is enforced after each pulse train, yielding a maximum behavioral event recording rate of 16.67 Hz for single-pulse events. At the maximum encoding of 39 pulses, the minimum interval between events extends to 99 ms (60 ms separation + 39 ms transmission), reducing the effective recording rate to approximately 10 Hz. Researchers should keep this in mind when designing paradigms centered on high-frequency behaviors, such as licking. Assigning longer pulse trains to frequently occurring events directly reduces the effective recording frequencies of the system.

An operational consideration is that this pulse train architecture creates vulnerability to event collisions: if a second Mod generates an event while a pulse train from the first Mod is still in progress, the overlapping pulses may be incorrectly decoded or lost entirely. Arduino controlled Mods do not suffer from this drawback as each Mod is blocking and prevents overlaps. However, the FED3 Mod is externally integrated and can overlap with Lickport and CS Mods. Event collisions are a concern only in multi-animal studies where two animals could independently activate different Mods (e.g., FED3 and Lickport) within the TTL transmission window. When two independent sources generate overlapping TTL pulse trains, the pulses can merge into a single extended train, producing either a mis-decoded (incorrect) event code or a decoding timeout. Across more than 50 real-world trials, such collisions were extremely rare (estimated (based on the absence of observed decoding errors across >50 trials, rather than systematic collision counting) at <0.1% of total events), primarily because typical inter-event intervals (>1 s) are long relative to the maximum pulse train duration (∼99 ms for a 39-pulse code). To minimize collisions, we recommend assigning shorter pulse trains to high-frequency events and avoiding simultaneous high-rate signaling from multiple independent modules or multiple animals on a single Core. Users are encouraged to stress-test their intended behavioral schedules for collisions prior to large-scale deployment.

#### 3.9.4 Serial Communication Reporting as Blocking in Control Loop

Arduino and FED3 control scripts depend on rapid loop execution to reliably sample sensor data, manage behavioral paradigms, and transmit events to the ESP32 logger. Serial print statements, commonly used during development for debugging and output reporting, introduce significant blocking behavior when left in production code. At 115200 baud, each Serial.println() call occupies the processor for several milliseconds while the string is transmitted, long enough to substantially reduce effective loop throughput and introduce lag into sensor sampling. This degrades the temporal fidelity of the system in proportion to how frequently such calls occur.

Researchers should ensure that all serial print statements are removed or disabled prior to protocol deployment to avoid missed operant events. In short, all serial debugging statements (e.g., Serial.println() calls) should be removed or disabled in production behavioral paradigms, as these blocking operations can degrade timing performance and violate the sub-millisecond latency guarantees. Serial calls that are guaranteed to be non-blocking (incur latency consequences) during mouse behavior (e.g., “Start” logging, “Start” and “Stop” json-html requests) are acceptable.

#### 3.9.5 NeuroHab Hz Recording Capabilities

As described above, the maximum behavioral event recording rate is 16.67 Hz, imposed by the combined constraints of pulse train length and the mandatory transmission separation delay.

While events may be transmitted at higher rates or in parallel, doing so substantially increases the probability of incorrect pulse decoding, and events may be lost due to pulse train overlapping or buffer overflow (Tanenbaum and Bos, 2014). For applications requiring higher recording frequencies, BNC input recording of external devices offers considerably greater throughput, supporting single-channel recording at up to 247 Hz (Figure 4a) for short pulse trains. Researchers synchronizing camera frame data with NeuroHab events should note that simultaneous pulse train delivery may result in frame loss, and this should be treated as a potential point of failure when such synchronization is critical to the dataset. Additionally, at high recording frequencies, buffer overflow may cause BNC2 input data to be written into BNC3 output files, or BNC3 input data into BNC4 output files, an artifact of the buffer memory layout (Tanenbaum and Bos, 2014) that should be monitored carefully. Figures 4a, 4b, 5a, and 5b illustrate the validated maximum recording frequencies achieved across single, dual, and triple channel configurations with n = 3 trials per condition and each trial consisting of between 807 to 7432 pulses. Cross-channel buffer artifacts have been observed empirically only under extreme, high-frequency stress conditions or from external EMF conditions. As a practical safe operating regime for multi-hour recordings with multiple active BNC channels, we recommend keeping sustained input rates below approximately 47.2 Hz per channel or explicitly increasing the logger’s internal buffer sizes and re-validating performance before deployment.

#### 3.9.6 RTC Module Limitations DS1307

The NeuroHab incorporates a DS1307 real-time clock (RTC) module for date-times of files and events alongside millisecond timestamps. The DS1307 is subject to drift, losing seconds to minutes of accuracy over multi-day operation. For precise synchronization, researchers should rely exclusively on the ESP32 microsecond-precision timestamp column (labeled “millis” and represents the time elapsed since session start) rather than the RTC-derived date-time values. If absolute wall-clock time is required (e.g., for circadian studies), we recommend replacing the DS1307 with a temperature-compensated RTC such as the DS3231 or disciplining timing to an external precision reference via a BNC input. For most behavioral-neural experiments, the relative millisecond timestamps are fully sufficient. In summary, the ESP32-derived microsecond-resolution relative timestamps are the authoritative time base for all fine-grained temporal analyses, including alignment to neural recordings; the DS1307 date-time field is provided only for coarse temporal context and file organization and should not be used for sub-second precision work due to drift.

#### 3.9.7 ESP32 Crystal Oscillation Imperfections and Impact on Accuracy

The ESP32 internal clock is governed by a crystal oscillator accurately within microseconds over short intervals. However, minor imperfections in crystal oscillation produce cumulative drift on the order of a few seconds per day (Espressif Systems, 2024b). In practice, this means that while millisecond timestamps are dependable across seconds and minutes of recording, the computed difference between an event at the start and an event at the end of a 24-hour session may deviate by several seconds from the true elapsed time. This should be considered a potential source of error in long, standalone recordings. When the NeuroHab is used in tandem with external data acquisition systems, however, drift becomes largely inconsequential, as temporal alignment is established relative to a shared reference. Relative synchronization between neural and operant timestamps therefore remains intact, only the absolute representation of elapsed time over extended intervals is affected. For most experimental applications this is a negligible concern; it becomes relevant only when microsecond precision must be maintained for analysis of intervals spanning hours or more, not alignment of events against cell activity.

#### 3.9.8 Memory Management for ESP32 Logging Designs

To sustain high recording rates, the NeuroHab ESP32 dedicates most of its onboard memory to logging buffers. Saving incoming events to buffers and then writing the buffers to disk in batches decreases missed events and frees the CPU for additional logic. Six buffers are allocated as 2D arrays, using uint32_t and uint64_t data types to store pulse counts and timestamps. Two parameters govern buffer behavior: buffer_size, which determines how many events can be held before a buffer switch is required, and numPins, which sets the number of active BNC recording channels. Increasing buffer_size supports higher recording frequencies but consumes more memory. Decreasing numPins frees memory for redistribution to buffer_size and other variables but reduces BNC channel availability. At the default values of numPins=4 and buffer_size=90, all three BNC channels are active with limited (but some) additional memory available for logic expansion. Users modifying these parameters should note that exceeding available memory will cause system crashes; guidance for proper memory management is available in the supporting GitHub documentation (Sun Lab, 2026).

#### 3.9.9 Lickport Module Sensitivity Limitations

Lickport activation relies on capacitive sensing, making proper mounting and calibration essential for reliable performance. Several factors influence the capacitance baseline and should be accounted for during setup. Lickports must not be mounted on metal surfaces where sensor contacts touch the mounting area, as this will produce false positive activations. Similarly, submerging or inconsistently soaking the contacts should be avoided, as inconsistent exposure introduces variance in the capacitance baseline. Activation thresholds must be set for rodent-scale capacitance, as human contact produces a larger capacitive signal due to differences in body size, erroneous activations from wire-to-surface contact indicate that the detection threshold is set too low. Finally, fluid within the delivery lines alters system capacitance relative to empty lines; calibration should therefore always be performed with the intended liquid in the plumbing. Calibration steps are available in the accompanying repository (Sun Lab, 2026). Lickport accuracy data is summarized in Table 2; the 96.83% fixed-trial accuracy (n=63 events) is attributed to camera frame-rate limitations at the detection boundary; concurrent electrical contact validation was not performed and represents a direction for future validation work.

#### 3.9.10 Mod Limitations

Operant Mods installed for behavior analysis in Home environments are limited by the response latency of sensors. The time between real event occurrence and sensor registration of the event (response latency) is unavoidable and variable across Mods. NeuroHab recording is less than 58 μs latency from sensor activation to log timestamp. Externally integrated Mods should be tested to confirm that response latency does not exceed 942 μs, calculated as 1000 μs (sub-millisecond requirement) minus the maximum combined logging latency of 58 μs (56 μs buffer switching + 2 μs ISR, Figures 12, 13), to ensure system latency remains below 1 ms. NeuroHab Mods (Lickports, Conditioned Stimulus, FED3) do not exceed this latency. Lickport capacitive sensors have a response latency of 223.48 μs (1 Lickport) to 615.88 μs (3 Lickports) (Figure 2), calculated from 200 samples per channel as the interval between Core loop start and confirmed activation, as sensors update once per loop. Abrupt activations that cross the detection threshold rapidly produce lower latencies, while gradual activations that approach the threshold slowly may produce higher latencies or fail to register entirely. Lickport latency and accuracy are dependent on threshold and sampling settings within the NeuroHab library and may require adjustment based on experimental requirements. FED3 IR sensors operate via interrupt service routines that set Boolean flags detected during the main loop. Because main loop duration varies with user programming, FED3 IR response latency is not fixed. Following NeuroHab modification, FED3 IR response latency does not exceed 60 μs, with a mean of 28.96 μs across 100 samples per port (Figure 8). Conditioned Stimulus modules exhibit negligible response latency, as they are non-blocking, activation-only devices with no sensor polling.

Response latencies for Lickports and FED3 devices were measured by recording the microsecond timestamps from the device architecture before sensor update to immediately prior to logging. The timestamps include the full loop execution to account for the operant paradigm execution time to represent the response latencies as accurately as possible. This loop execution time is seen in the even distribution of the FED3 latency (Figure 8). The shortest response latencies result from ISR returning to just before sensor evaluation, conversely, the longest latencies required the full completion of the next loop to elapse. Lickport response latencies were measured similarly, with the initial timestamp saved before capacitive sensor update and elapsed time reported after sensor registration. The consistent distribution of the Lickport response latencies results from the lack of an ISR, so the program executes the same set of instructions each loop (Figure 2).

#### 3.9.11 Solenoid Back EMF Handling

For the delivery of precise fluid volumes via Lickports, 12V DC solenoid valves are controlled through relay modules and digital signaling pins. As solenoid valves open and close, the collapsing electromagnetic field generates back-EMF across connected circuits, which can erroneously trigger TTL pulse detection and damage downstream components. To suppress this, UF4007 flyback diodes are wired from the normally open relay terminal to ground for each solenoid circuit. The UF4007 was selected for its fast recovery time and high resistance to reverse current (Horowitz and Hill, 2015).

### 3.10 System Cost

The NeuroHab, including all modules, requires approximately $1,400 in materials including bulk parts useable in future builds. The FED3 Mod and LeeCo solenoid valves account for 61% of the total build cost (∼$860 combined) representing a majority cost driver for the full system.

Excluding the FED3 and supplementing US Solid Valves in place of the more expensive LeeCo valves reduces the cost to build an additional system from leftover components to only $170 and only $106 if a new Home is not constructed. After purchasing a second FED3 and using LeeCo valves, the cost for a second complete system is $995 of which the FED3 and LeeCo valves account for approximately 86% of the total cost. Depending on operant requirements, liquid delivery and CS systems can be set up for only $170 per unit. This includes 3 Lickports and 3 CS modules which can be installed in 3 mouse cages to reduce the cost of the set up to only $56 per cage. By comparison, the Med Associates single-chamber operant conditioning package comprising a Behavioral Classic Chamber with nose poke and USB operating software for up to eight chambers is quoted at $12,952 before shipping with the cage itself costing $3,993 (Med Associates Inc., 2026). This represents a conservative 9-fold cost reduction for one system and 4.7 times cheaper for 8 systems ($41,225 vs $8,762). Further, large setups which do not rely on FED3 feeding can achieve far greater cost savings at scale. Generally, cost is directly associated with the complexity of the operant setup, bounded by the figures above, existing lab resources, and researcher time. Note that the NeuroHab cost figure excludes assembly labor and 3D printing time, whereas the Med Associates quote includes assembled hardware; additionally, the Mini two-photon microscope (∼$10,000+) is a separate cost not included in the NeuroHab figure.

## 4. Discussion

### 4.1 Overview

The NeuroHab was developed to address a key challenge in operant behavioral neuroscience: cost-effective access to a system with high temporal resolution for synchronization with neural recording modalities such that neural encoding and decoding of operant tasks can be conducted. Collectively, these barriers have historically forced laboratories to invest significant time and resources building bespoke rigs for each new experiment, often arriving at solutions that are difficult to validate, reproduce, or extend.

The NeuroHab addresses these challenges through a modular, open-source architecture that standardizes common behavioral components while preserving the flexibility to design and execute novel operant paradigms. Existing open-source behavioral systems including pyControl and custom Arduino-based platforms have established the value of open, customizable tools; the NeuroHab extends this paradigm by integrating <1 ms event logging with home-cage compatible hardware and explicit two-photon imaging synchronization in a single unified system. By integrating event logging, stimulus delivery, and reward dispensing within a unified control system, the NeuroHab reduces redundant development effort and accelerates experimental throughput in ways that neither commercial nor traditional custom solutions readily support. Researchers retain full programmatic control over paradigm design without engaging in low-level system engineering (Espressif Systems, 2024a; Tanenbaum and Bos, 2014), and the system’s compatibility with external neural recording hardware removes a significant barrier to multimodal experimental design.

From a cost standpoint, the NeuroHab represents a substantial reduction relative to commercial alternatives. Unlike commercial systems, which are typically designed for fixed experimental paradigms and rarely support concurrent neural data acquisition, the NeuroHab is explicitly engineered for integration with electrophysiology and calcium imaging hardware, enabling temporally precise multimodal studies that would otherwise require substantial additional infrastructure.

### 4.2 Capacitive Sensor Application for Broad Use Cases

The capacitive sensors used for Lickport activation are not inherently limited to fluid delivery applications. Because these sensors measure the capacitance of any connected conductive object, they can be attached to arbitrary conductive surfaces to detect and record animal contact across a wide range of experimental contexts. This flexibility extends the utility of the NeuroHab well beyond standard licking paradigms. For example, a set of metal floor pads wired to Core terminals could detect positional preference or zone occupancy within the home cage, with conditioned stimuli or rewards delivered contingent on pad activation to probe area preference, place aversion, or disruption of spatial behavioral patterns. The range of applicable designs is broad, and the low latency and precision of our architecture make it well suited for operant paradigms evaluating neural activity which require reliable contact detection across novel surface configurations.

### 4.3 Applications for Multiple Animal Studies in Separate and Shared Environments

The NeuroHab architecture is well suited for extension to multiple animal operant studies. In spatially separated configurations, each animal is assigned a dedicated Home environment with its own Mod installations, and event logs map directly to individual animals without ambiguity. In shared environments, logs record sensor activation without identifying the activating individual. While useful for integrating the required operant task or conditioned stimulus, the researcher is left to design per animal identification protocols. Theoretically, RFID chips may be utilized along with external sensors to evaluate specific individuals’ movement within their environment in tandem with Mod activation to identify individual behavior from a group.

The primary hardware consideration for multi-animal studies is the potential for simultaneous Mod activations to produce overlapping TTL pulse transmission, which could result in incorrect event decoding at the Core. This risk is mitigated by the program structure of the NeuroHab: simultaneous activations of Lickport or CS modules cannot occur by design, limiting race conditions (Tanenbaum and Bos, 2014) from FED3 and Mod events. Given the millisecond transmission speeds of the Core, such collisions would require near-perfect overlap between independent activations and are therefore expected to be rare in practice. However, researchers designing multi-animal paradigms should review the Hz recording capabilities and TTL communication limitations outlined in Sections 3.9.3 and 3.9.5 to ensure their experimental design remains within validated performance bounds and should consider that collision risk increases with the number of independent event sources.

### 4.4 Additional Mod Integration via Sensor Libraries

The NeuroHab Core exposes the unused digital and PWM pins on the onboard Arduino, providing substantial capacity for researcher-defined hardware expansion. The broad compatibility of Arduino architecture with open-source sensor libraries enables straightforward integration of additional Mods without modification to the Core logging or control structure. Environmental sensors such as humidity sensors, distance detectors, motion sensors, servo actuators, and custom lighting systems are among the many hardware categories compatible with this framework. Each can be incorporated into operant paradigm design while leveraging the existing event recording and synchronization capabilities of the NeuroHab Core, lowering the barrier to novel experimental configurations that would otherwise require independent rig development.

### 4.5 Closing

The NeuroHab represents a practical and accessible solution to infrastructure challenges in behavioral neuroscience. By integrating precise event logging, modular reward and stimulus delivery, and compatibility with neural recording hardware into a single open-source platform, it reduces the time, cost, and technical expertise required to conduct high-fidelity operant research. The system’s <1 ms behavioral event logging latency, with microsecond-resolution timestamps and millisecond-scale synchronization to external neural recording hardware, combined with its flexible and extensible architecture, positions it as a foundation upon which diverse experimental paradigms can be built and refined without redundant engineering effort. As neuroscience increasingly demands multimodal approaches that bridge behavior and neural activity, tools that lower the barrier to temporally precise, reproducible, and combinable experimental designs become essential. The NeuroHab is offered as a contribution toward that goal. Software and hardware schematics are available in the accompanying GitHub repository (Sun Lab, 2026). Plans for commercial availability are currently underway, with production details to be finalized following the release of this paper.

**Table 3.**
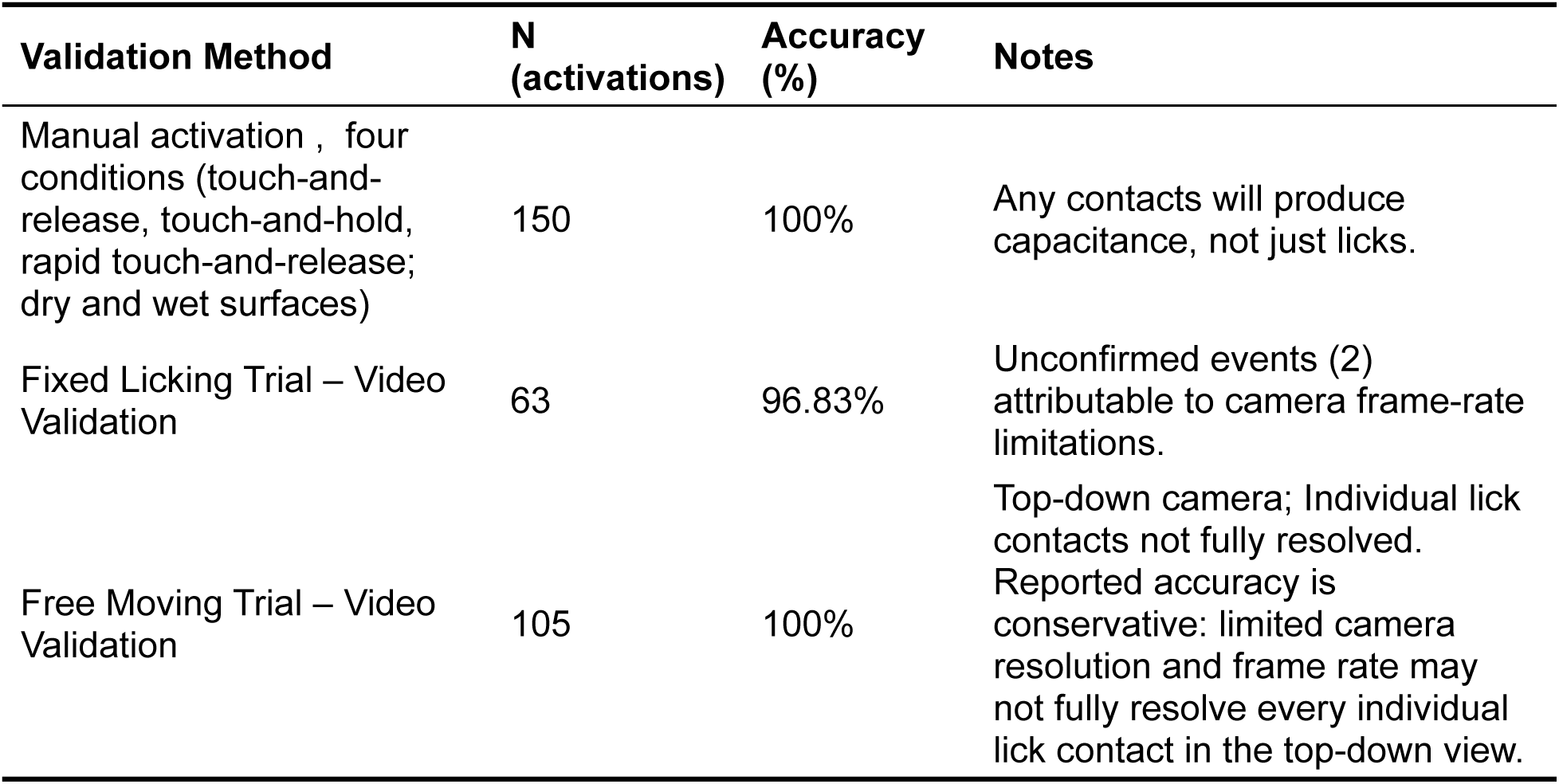
Lickport validation trials.

## Author contributions

Q.-Q.S. designed the experiments, drafted the manuscript, and acquired funding. S.C. co-designed the project, developed the electronics and firmware, manufactured the hardware, and performed the validation experiments. R.J. measured animal operant behavior, provided feedback on earlier versions of the hardware, and edited the manuscript.

## Data Availability

All hardware schematics, firmware (Arduino and ESP32 code), and analysis scripts are available in a public GitHub repository: https://github.com/wyomingneuron/NeuroHab.

Representative sample log files from key experiments, including 16-hour home-cage sessions, licking validation, and two-photon synchronization tests, have been deposited in a /sample data directory within the repository. Additional raw data are available from the corresponding author upon reasonable request.

## Acknowledgements

The authors thank the Kravitz Laboratory at Washington University School of Medicine for the development of the FED3 platform upon which the food delivery integration described herein is based. We thank Dr. Nuo Li, Duke University, for providing the drawings and circuit board for the initial version of the Lickport, which we fully redesigned in this manuscript. This work was supported by grants from the National Institute of Mental Health (R21MH131363, R21MH141703) and the National Institute of General Medical Sciences (2P20GM121310) and the Office Of The Director, National Institutes Of Health (OD).

## References

1. Akam T, Lustig A, Rowland JM, Kapanaiah SKT, Esteve-Agraz J, Panniello M, Márton CD, Kätzel D, Bhatt DL, Bhatt D (2022) Open-source, Python-based, hardware and software for controlling behavioural neuroscience experiments. eLife 11:e67846.

2. Arduino (2023a) Arduino MEGA 2560 Rev3. Available at https://docs.arduino.cc/hardware/mega-2560/.

3. Arduino (2023b) Serial communication documentation. Available at https://docs.arduino.cc/learn/communication/serial.

4. Badger P, Stoffregen P (n.d.) CapacitiveSensor library. Available at https://github.com/PaulStoffregen/CapacitiveSensor.

5. Buzsáki G (2004) Large-scale recording of neuronal ensembles. Nat Neurosci 7:446–451.

6. Crawley JN (2012) What’s wrong with my mouse? Behavioral phenotyping of transgenic and knockout mice, 2nd ed. Hoboken, NJ: Wiley.

7. De La Crompe B, Schneck M, Steenbergen F, Schneider A, Diester I (2023) FreiBox: a versatile open-source behavioral setup for investigating the neuronal correlates of behavioral flexibility via 1-photon imaging in freely moving mice. eNeuro 10:ENEURO.0469-22.2023.

8. Espressif Systems (2024a) ESP-IDF FreeRTOS (SMP) changes. Available at https://docs.espressif.com/projects/esp-idf/en/latest/esp32/api-guides/freertos-smp.html.

9. Espressif Systems (2024b) ESP32-DevKitC V4 user guide. Available at https://docs.espressif.com/projects/esp-dev-kits/en/latest/esp32/esp32-devkitc/user_guide.html.

10. Horowitz P, Hill W (2015) The art of electronics, 3rd ed. Cambridge, UK: Cambridge University Press.

11. Kapanaiah SKT, Strahnen D, Akam T, Bannerman DM, Kätzel D (2021) A low-cost open-source 5-choice operant box system optimized for electrophysiology and optophysiology in mice. Sci Rep 11:22776.

12. Mathis A, Mamidanna P, Cury KM, et al. (2018) DeepLabCut: markerless pose estimation of user-defined body parts with deep learning. Nat Neurosci 21:1281–1289.

13. Matikainen-Ankney BA, Earnest T, Ali M, Casey E, Wang JG, Sutton AK, Legaria AA, Lotun A, Bhatt D, Nguyen KP, Keiflin R, Kravitz AV (2021) An open-source device for measuring food intake and operant behavior in rodent home-cages. eLife 10:e66173.

14. Med Associates Inc. (2026) Quote MQ00054566: behavioral classic chamber package with nose poke for mouse (MED-307A-B2), USB operating package for up to 8 chambers (MED-SYST-8-USB), power cable SG-210CP-25. Issued March 26, 2026. Total: USD $12,952.00.

15. Nguyen KP, O’Neal TJ, Bolonduro OA, White E, Kravitz AV (2016) Feeding Experimentation Device (FED): a flexible open-source device for measuring feeding behavior. J Neurosci Methods 267:108–114.

16. Qian L, Liu Y, Chen Y, et al. (2026) High-throughput two-photon volumetric brain imaging in freely moving mice. Nat Commun 17:206.

17. Sun Lab (2026) NeuroHab. GitHub. Available at https://github.com/wyomingneuron/NeuroHab.

18. Tanenbaum AS, Bos H (2014) Modern operating systems, 4th ed. Upper Saddle River, NJ: Pearson.

19. Zong W, Wu R, Li M, Hu Y, Li Y, Li J, Rong H, Wu H, Xu Y, Lu Y, Jia H, Fan M, Zhou Z, Zhang Y, Wang A, Chen L, Cheng H (2017) Fast high-resolution miniature two-photon microscopy for brain imaging in freely behaving mice. Nat Methods 14:713–719.

